# Origin of the bidirectionality of cerebrospinal fluid flow and impact on long-range transport between brain and spinal cord

**DOI:** 10.1101/627166

**Authors:** Olivier Thouvenin, Ludovic Keiser, Yasmine Cantaut-Belarif, Martin Carbo-Tano, Frederik Verweij, Nathalie Jurisch-Yaksi, Pierre-Luc Bardet, Guillaume Van Niel, François Gallaire, Claire Wyart

## Abstract

The circulation of cerebrospinal fluid (CSF) plays pivotal roles for body axis formation and brain development. During embryogenesis, CSF is rich in particles and proteins and flows bidirectionally in the central canal. The origins of bidirectional flow and its impact on development are unknown. Experiments combined with modeling and simulations demonstrate that the bidirectionality of CSF flow is generated locally by caudally-polarized motile cilia confined to the ventral wall of the central canal. Such active bidirectional flow of the CSF accelerates the long-range transport of particles propagating rostrally and caudally. In addition, spontaneous muscle contractions increase local CSF flow and consequently enhance long-range transport of extracellular lipidic particles. Focal ablation of the channel connecting brain ventricles to the central canal reduces embryo length, indicating that long-range transport contributes to embryonic growth. Our study also demonstrates that at this early stage, motile cilia ensure the proper formation of the central canal.

## Introduction

Precise control of flow in biological systems is essential to transport critical molecules to the different cells in the organism. In order to respond to the cells demands, passive diffusion in tissues imposes that every cell in the organism is close enough to a supply source. For organisms larger than a few hundreds of microns, diffusion alone is too slow to achieve supply of nutrients to all cells in the body. Therefore, organisms have developed a large variety of flows to accelerate and orient the transport of key molecules. Blood flow is the main carrier of glucose and oxygen in most organisms and is mostly driven by the pressure gradient caused by the heart pump^1^. Another large category of biological flows, including intestine, esophageal or lymph flows, relies on peristalsis, where muscles contractions drive the fluid in the direction of propagation of the contraction wave^2–6^. Another major class of fluid flows relies on the asymmetric beating of motile cilia or flagella, which are efficient motors to generate directional flows at small scales and to mix fluids in biological systems^7–10^. For instance, cilia-driven flows were demonstrated to be critical for left/right asymmetry in the developing embryos of many species including humans^11^, in the Kupffer’s vesicle in zebrafish^10,12^, and in the node in mammals^13–15^. Cilia-driven flows can also be observed in the pronephros^16,17^, in the respiratory tract ^18^, and in the nasal cavity^19^. The motility of cilia in brain ventricles is also correlated with the cerebrospinal fluid (CSF) flow in zebrafish^17,20,21^, African clawed frog^22,23^ and rodents^24^.

Understanding CSF circulation throughout the body is crucial as this biological fluid plays numerous roles in the development of brain and spinal cord. The CSF provides the hydromechanical protection of the brain^25^, enables the transport of nutrients and waste^25,26^, exosomes^27–31^, and signaling molecules impacting neurogenesis and neuronal migration in the brain^32–35^. The CSF circulation has recently been linked to body axis formation in the embryo and spine morphogenesis in juveniles/adults^21,36,37^. During embryogenesis, mutations affecting ciliogenesis, cilia polarity or motility lead to defects of transport along the rostrocaudal axis and body axis formation^36,38^.

Historically, it has been generally assumed that CSF flow along the walls of all cavities must be driven by motile cilia. In the brain ventricles, the direction of beating cilia correlates indeed with the direction of CSF flow^20,24^. However, in the long cylinder of the central canal, the dynamics of CSF flow is strikingly different than in the brain ventricles. We recently showed in 24 hours post fertilization (hpf) zebrafish embryos that the CSF flows bidirectionally^31,39^. To our knowledge, these observations report for the first time a constant bidirectional flow in a single channel in biological systems. Interestingly, body-axis defects observed in mutants with defective cilia appear by 30 hpf, i.e. few hours before cilia paving the walls of the brain ventricles become motile and a directed CSF flow emerges in the brain ventricles^20,40^. It suggests that proper dynamics of CSF flow in the central canal, not in the brain ventricles, is critical for embryonic body axis formation.

Since the mechanisms leading to the bidirectionality of CSF flow in the central canal were unknown, we developed an automated method for quantifying CSF flow in the central canal of zebrafish embryos. We show that motile cilia are critical to form the lumen of the central canal, as the canal collapses in mutants with defective cilia. We combined experimental, numerical and theoretical approaches to test the role of the spatially-asymmetric distribution of motile cilia in the generation of a bidirectional flow. We developed a general and simple hydrodynamic model to account for the contribution of cilia, and show that this model can be applied as a tool in many cilia-driven flows in confined environments. Additionally, we show experimental and theoretical evidence for an intricate relation between local flow and long-range transport of particles in the CSF. We demonstrate that this bidirectional CSF flow in the central canal is enhanced by muscle contractions, which accelerate bidirectional transport of particles both rostrally and caudally. We show that the CSF flows unidirectionally from the diencephalic/mesencephalic ventricle to the entrance of the central canal via a long and thin funnel. *In vivo* two-photon ablation of this funnel leads to a reduction in growth, suggesting that the exchange of critical particles between brain ventricles and central canal is necessary for optimal embryogenesis.

## Results

### Quantification of CSF flow in the central canal

Understanding the mechanisms determining CSF flow in the central canal requires an *in vivo* quantification and a theoretical framework model with minimal parameters to explain it. To establish quantitative and reproducible flow profiles along the dorsoventral (D-V) axis of the central canal (CC), we develop an improved automated analysis inspired by our previous manual kymograph measurements^31,39^. By co-injecting 3,000 molecular weight (MW) red dextran with 20 nm-diameter green fluorescent beads in the diencephalic/mesencephalic ventricle (DV), we measure the volume of cavities and particle trajectories.

The dextran rapidly diffuses over all fluid-filled cavities, including the CC (**Figure 1a**), enabling its visualization and the assessment of the injection quality. Fluorescent nanobeads show random Brownian motion superimposed to a smooth displacement along the streamlines (**Supplementary Movie S1**). Bead trajectories reveal the bidirectionality of CSF flow along the rostrocaudal axis and the presence of a few regions of recirculation, commonly referred to as vortices (**Figure 1b**). The flow analysis relies on the permutation of the 3-D (2-D in space and time) stack depicting trajectories of fluorescent beads (**Figure 1c1**) to form kymographs at various D-V positions in the CC (**Figure 1 c2, c3, c4, and Supplementary Movie S2**). The typical velocity profile is bimodal with rostro-caudal flow in the ventral side and caudo-rostral flow in the dorsal side, and reverses close to the center of the CC (normalized D-V position = 0.53, **Figure 1d** from 110 wild type (WT) embryos each imaged in 2 rostro-caudal positions). Automated kymographs show better reproducibility compared to particle tracking velocimetry (PTV) or particle imaging velocimetry (PIV), as discussed in Methods section. Velocity profiles show ventrally a maximal speed of 4.78 ± 0.79 μm.s^-1^ (D-V = 0.29 on average) and dorsally a minimal speed of −4.80 ± 0.82 μm.s^-1^ (D-V = 0.82 on average) (**Figure 1d1 and 1d2**) and is nearly symmetrical (D-V =0.53 on average). Similarly, the average speed throughout the CC is zero (−0.02 ± 0.15 μm.s^-1^), refuting the hypothesis of a global pressure gradient as the unique driver of the flow in the CC.

**Figure 1.**
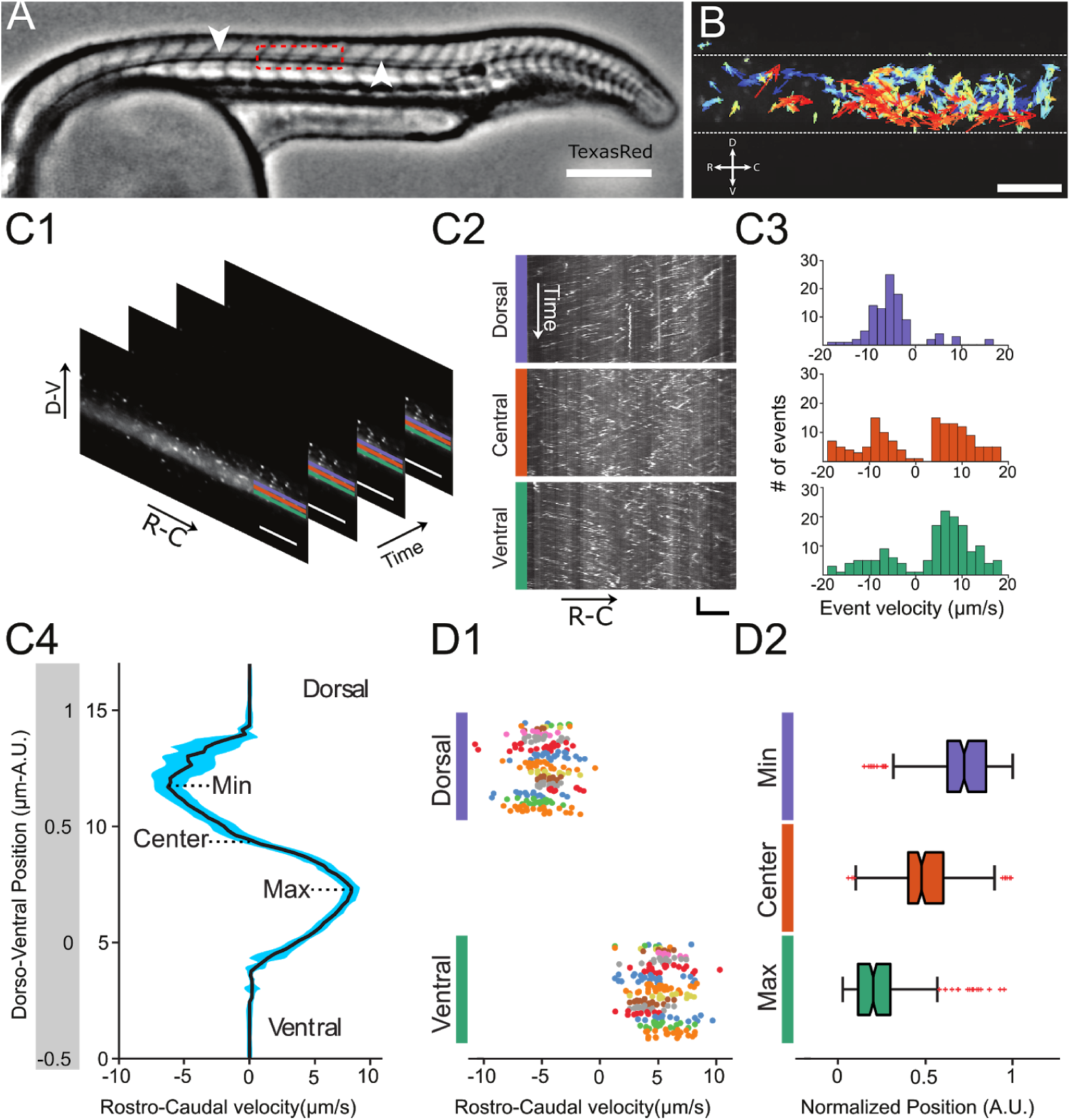
Automated CSF flow analysis in the central canal reveals a bidirectional velocity profile. (A) Inverted contrast widefield image showing a 30 hpf zebrafish embryo injected with Texas Red: Texas Red fills all fluid-containing cavities, including ventricles, central canal (CC, white arrows), floor plate, somite boundaries, and blood vessels. (B) 20 nm fluorescent particles instantaneous velocities, as measured with a particle tracking velocimetry (PTV) algorithm, inside the central canal in a region corresponding to the red box in (A). The length, direction, and color of all arrows are coded according to the instantaneous bead velocity, ranging from −10 (blue) to 10 μm/s (red). (C1) Time series showing subsequent images of fluorescent beads injected in the DV and transported down the CC used to generate kymographs at different dorso-ventral (D-V) positions. (C2) Example kymographs computed in a dorsal, a central, or a ventral position in the CC. Each line represents the trajectory of one bead, projected along the rostro-caudal (R-C) axis. The slope of each trajectory gives a bead velocity projected onto the R-C axis. (C3) Histogram of velocities obtained at the 3 positions shown in (C2). (C4) Velocity profile (mean ± s.e.m. in blue) calculated for all D-V positions in the CC for one WT embryo showing a maximum in the ventral side and a minimum in the dorsal side. (D1) Values of minimal (extremum in the dorsal part) and maximal velocities measured on 110 WT zebrafish embryos. Each dot represents the extremal values of one profile, and each color presents an experiment performed on siblings. 2 R-C positions are sampled per embryo. Extremal velocities are respectively 4.78 ± 0.79 μm.s^-1^ (ventral CC) and 4.80 ± 0.82 μm.s^-1^ (dorsal CC). (D2) Values of normalized D-V position of minimal, null, and maximal speed in the dorsal, center and ventral position in the CC (median positions are respectively 0.82, 0.53, and 0.29). Horizontal scale bar is 15 μm in (a, b), and vertical scale bar is 5 s in (b).

### Central canal geometry and properties of the motile cilia

To build a theoretical model of CSF flow, we quantify essential parameters such as the canal geometry and the local cilia dynamics. The central canal geometry, as estimated by Dextran injections, corresponds to a cylinder (**Figure 2a1, 2a2**) of diameter 8.9 ± 0.9 μm (129 WT embryos, **Figure 2a3**) with a very small dorsal extension showing low fluorescence level in the uppermost position, as previously observed with antibody staining for tight-junctions^41^. The dynamics of motile cilia in the CC are investigated in 30 hpf *Tg(β-actin:Arl13b-GFP)* embryos in which GFP labels all cilia^31,39,42^. Density and motility of cilia are higher in the ventral CC as previously reported^31,39,42^. Movies recording the cilia beating illustrate the high density of cilia and diversity in ciliary length and beating frequency (**Figure 2b**, **Supplementary Movie S3** and **Supplementary Figure S1**). While some cilia beat at high frequency (at least 45 Hz, see **Methods**) with a single frequency pattern, others beat at lower frequency (∼13 Hz) with more complex and irregular patterns. An automated quantification of cilia beating frequency using temporal Fourier transform on each pixel (**Figure 2b1**) reveals distinct spots of constant frequencies (**Figure 2b2**), likely corresponding to the displacement of a single cilium. For each spot, we extract the main frequency, orientation, length and height (**Figure 2c1-4**), indicating that on average cilia beat at 37.3 Hz with a caudal tilt of 65.0° towards the tail (median of the absolute value, see **Methods**), and are 6.3 μm long, thereby occupying a 2.9 μm height in the ventral canal, representing a bit less than half of the central canal. The caudal tilt is consistent with previous reports^39,42^. The measured frequencies are higher than the 12-15 Hz previously reported in zebrafish central canal^17^ (see **Methods** explaining the discrepancies), but are consistent with beating frequencies of ependymal cilia located in the brain ventricles of rats^43^ and zebrafish larvae ^20,43^.

**Figure 2.**
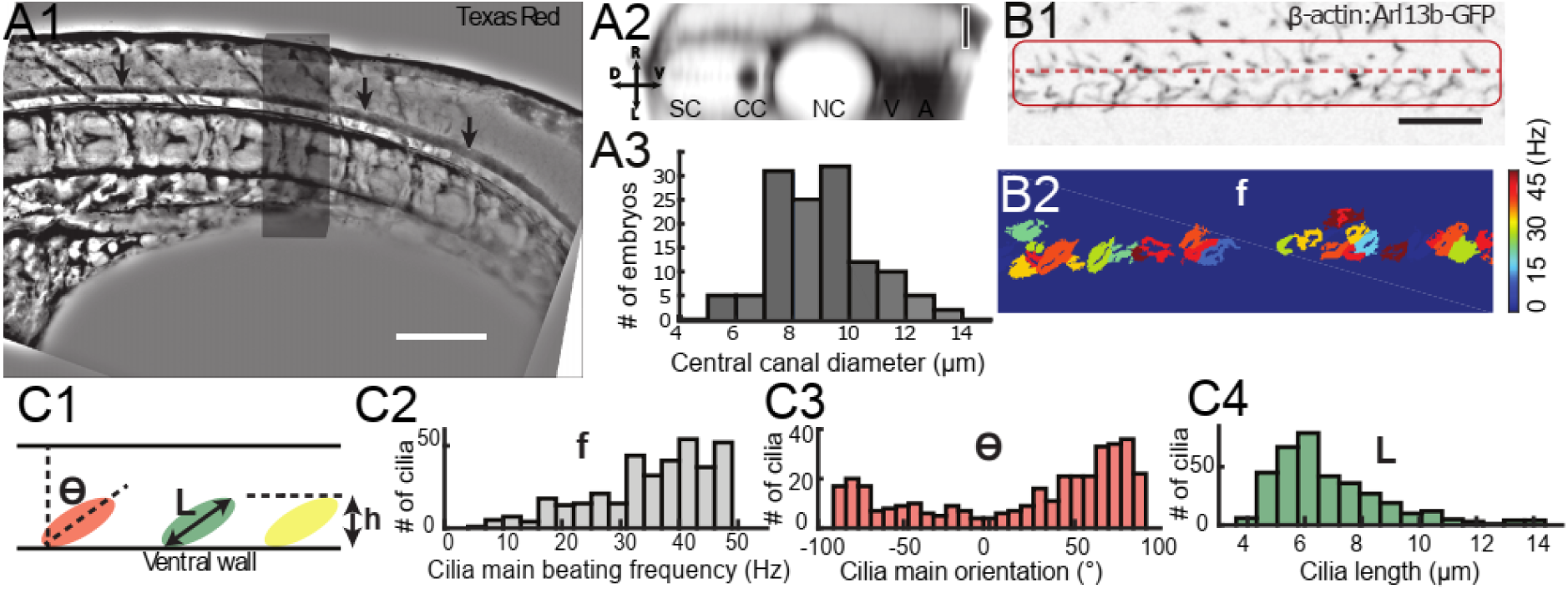
Geometry of the central canal and properties of motile cilia. (A1, A2) Central canal filled with Texas Red injected in the brain ventricles shown laterally from maximal Z-projection stack of 75 inverted contrast images acquired with 1 μm Z-step (A1) and corresponding vertical profile (A2, 5x rescale on the axial dimension enabled similar pixel size in X/Y and Z): SC, spinal cord (light grey), CC, central canal (black, and black arrows in (a1)), NC, notochord (white), V, vein and A, artery (black). (A3) Histogram of central canal diameter from N= 129 WT embryos, with a median value of 8.9 ± 0.9 μm (mean ± 0.5 s.t.d.). (B1, B2) Lateral view in the CC where cilia are labeled with GFP in *Tg (β-actin: Arl13b-GFP)* transgenic embryos (B1) enables to draw map of local main beating frequency (B2) from the fluorescence signal obtained by calculating the fast Fourier transform of each pixel temporal trace. Regions of constant frequency are color-coded and supposedly correspond to single cilium beating. (C1) Scheme presenting some parameters of interest that will be extracted from regions of constant frequency obtained in (B2) and fitted as ellipses to obtain their mean orientation θ, their major axis (L), associated to the cilia maximal length, and their projection along the D-V axis, which instructs on the portion of central canal occupied by beating cilia. (C2, C3, C4) Histograms of parameters extracted in many regions of constant frequency of N=22 β-actin: Arl13b-GFP 30 hpf embryos, for a total of 360 cilia analyzed in total. A central frequency of 37.3 Hz (median frequency with s.d 10.0 Hz) (C2), a mean absolute orientation |θ| 65.0° (s.d. 23.4°) (c3), a length L of 6.3 μm (s.d. 2.0 μm) (C4) as well as a beating height of 2.9 μm (s.d. 1.9 μm) (not shown) are found. Scale bar is 50 μm in (A1), and 15 μm in (A2, B1, B2).

### The asymmetric distribution of beating cilia generates a bidirectional flow

The CC can be considered as a cylinder (**Figure 2a**) containing numerous motile cilia along the ventral wall (**Figure 2b**). Modeling the effect of each individual cilium beating would be extremely challenging. Since the CSF bidirectional flow does not vary over time and over rostro-caudal axis (**Figure 1c4**), we build a two-dimensional model that homogenizes cilia contribution as a constant force. The model divides the CC in two regions of equal thickness h = d/2 (**Figure 3a1, 3a2**). The dorsal region contains passive cilia whose influence is neglected, while active cilia generate a constant volume force f_v_ within the ventral region. Dimensional analysis gives f_v_ =αμf/h, where f is the cilia mean beating frequency (40 Hz), μ is the CSF dynamic viscosity and α is a numerical factor of order unity. The Stokes equation controls flow at small scales as inertial terms are negligible in comparison to viscous terms, and simplifies in this simple 2-D geometry to yield the velocity profiles in both regions:

**Figure 3.**
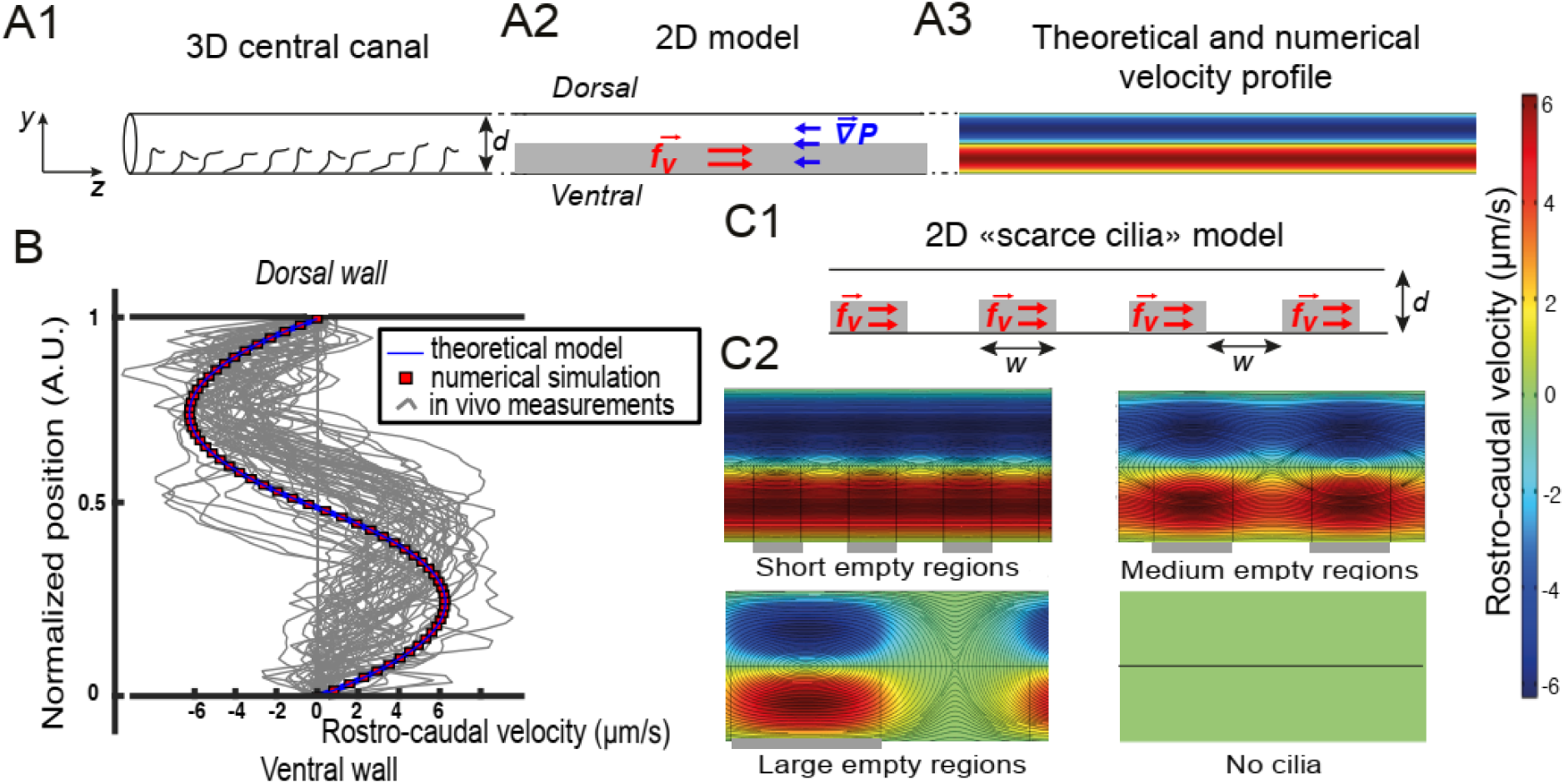
Theoretical and numerical results explain how motile cilia generate local CSF flow. (A1) Schematic of the three-dimensional central canal that can be reduced to a quasi-cylinder whose ventral wall is paved by regularly positioned motile cilia, and whose dorsal wall is composed of more loosely distributed passive cilia. (A2) The model reduces the geometry to a bidimensional channel of thickness d, composed of two main regions occupying the ventral and dorsal halves of the channel. In the ventral region, the cilia contribution is modelled as a constant bulk force. A constant gradient of pressure, present in the whole central canal, opposes the cilia-induced force and ensures a global zero flux on a cross section. (A3) Numerically and theoretically calculated velocity maps, where dark blue represents highest rostral velocities (negative values), while dark red holds for the highest caudal velocities (positive values). (B) Numerical (red squares) and theoretical (blue line) velocity profiles, composed of two parabolas matched at the center of the cross-section (normalized position = 0.5). They are convincingly compared to a superimposition of *in vivo* measurements obtained on 57 WT siblings. (C1) “Scarce cilia” model, accounting for the spatially inhomogeneous defects in cilia structure or motility (as observed in some mutants with defective cilia). The model reduces these defects to regions in the ventral side that are fully passive and where no force applies. Numerical simulations enable to derive the flow with an arbitrary distribution of these passive regions. (C2) Maps of characteristic streamlines obtained for various aspect ratios w/d of the passive regions. Small passive regions show most of the streamlines parallel to the walls. A central recirculation, or vortex is gradually more present in between cilia regions as the lateral extension of the passive region increases. The corresponding distribution of pressure in the central canal for these different cases is represented in **Supplementary Figure S2**.

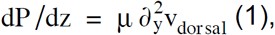

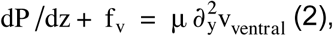

Where dP /dz is the common pressure gradient applied to both regions, and 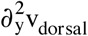 is the second y-derivative of the rostro-caudal velocity v_dorsal_(y). Matching velocity and shear rate at the boundary between both regions (y = h), the derived flow obeys to two parabolic profiles:

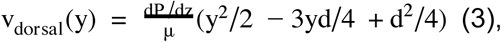

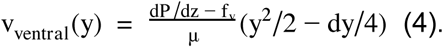

To solve this system of equations, one needs a third relation given by the constraint of zero flux across the diameter of the central canal as observed *in vivo*:

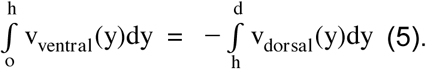

With a unique fitting parameter α chosen as 0.5, (leading to f_v_= 4000 N/m^3^) we obtain a symmetric bimodal flow with extreme velocities equal to ±5 μm/s (**Figure 3a3**), and an axial pressure gradient dP /dz = 2000 N/m^3^. We additionally solved the complete Stokes equation numerically, using Finite-Elements Methods simulations (COMSOL software) to confirm the hypotheses of the theoretical model. Theoretical and numerical predictions are in good agreement with 57 velocity profiles measured *in vivo* (**Figure 3b**).

The model is further refined to account for the spatial inhomogeneity in the cilia distribution in the ventral region, that we design as spots of constant volume force f_v_ regularly distributed along the ventral wall of CC with a period w (**Figure 3c1**). With this “scarce cilia” model, numerical simulations show that recirculation regions appear between these spots (**Figure 3c2**). The amplitude of the recirculation regions is more pronounced as w increases. For large passive regions of width larger than the diameter of the CC (w > d), all the streamlines are located within the vortices. Counter-intuitively, this model predicts that vortices may originate not from cilia dynamics, but rather from the local absence of motile cilia in the ventral side on a distance larger than d. Note that the presence of passive regions decreases the pressure in the CC proportionally to the fraction of passive regions (**Supplementary Figure S2a**). We can further generalize these results by showing that differences in beating frequency between active neighboring cilia lead to the emergence of recirculation regions (**Supplementary Figure S2d**).

### Cilia activity controls CSF long-range transport, CC structure, and local flow

To verify that asymmetric cilia beating is solely responsible for the bidirectional flow i*n vivo*, we injected beads in the diencephalic ventricle of several mutants for which either ciliogenesis or ciliary motility is affected. Consistently with previous reports^36,39^, long-range transport of beads down the central canal is altered in mutants with defective cilia (**Figure 4a1, 4a2**). In paralyzed mutant embryos several hours post injection, no particles entered the CC, as shown here in *traf3ip*^*tp49d/tp49*^, commonly referred to as *elipsa*^*^44^*^. Besides, we notice that the CC geometry is dramatically altered in *elipsa* as well as in *dnaaf1*^*tm317b/tm317b*^, (referred to as *lrrc50*^*^45^*^), in the newly-generated mutant allele *foxj1a*^*nw2*^ (see **Methods** and characterization in **Supplementary Figure S3**), and in *cfap298*^*tm304/304*^, referred to as *kurly*^*^46^*^ (**Figure 4b,c**). While *lrrc50* embryos show complete loss of cilia motility^45^, *foxj1a*^*nw2*^ mutants retain normal primary cilia but show a decreased density of glutamylated cilia in the central canal (**Supplementary Figure S3**). Glutamylated cilia in the CC of WT embryos are caudally-polarized and ventrally-located and most likely correspond to motile cilia (**Supplementary Figure S3 e1,e2**). The severe reduction of glutamylated cilia *foxj1a*^*nw2*^ mutants is consistent with the previous report showing the necessity of *foxj1* genes for formation of motile cilia^55^. In the mutant *elipsa*^39,44^, cilia partially degenerate. The mutant allele *cfap*^*tm304*^ is characterized by a thermosensitive loss of cilia motility^46^. In all four mutants with defective cilia that we tested here, the CC forms but is flattened along the D-V axis and tortuous compared to WT siblings (**Figure 4b1, b2, 4c)**. In contrast, we measure no difference in the CC diameter in the mutant *scospondin*^*icm15/icm15*^, which also exhibits a curled down phenotype but with normal cilia properties and activity^39^, suggesting that cilia integrity and motility is a critical parameter to open the CC. It also suggests that previous observations describing defects of transport in mutants with defective cilia^36^ could be simply explained by the collapse of the CC, and not only by CSF flow defects.

**Figure 4.**
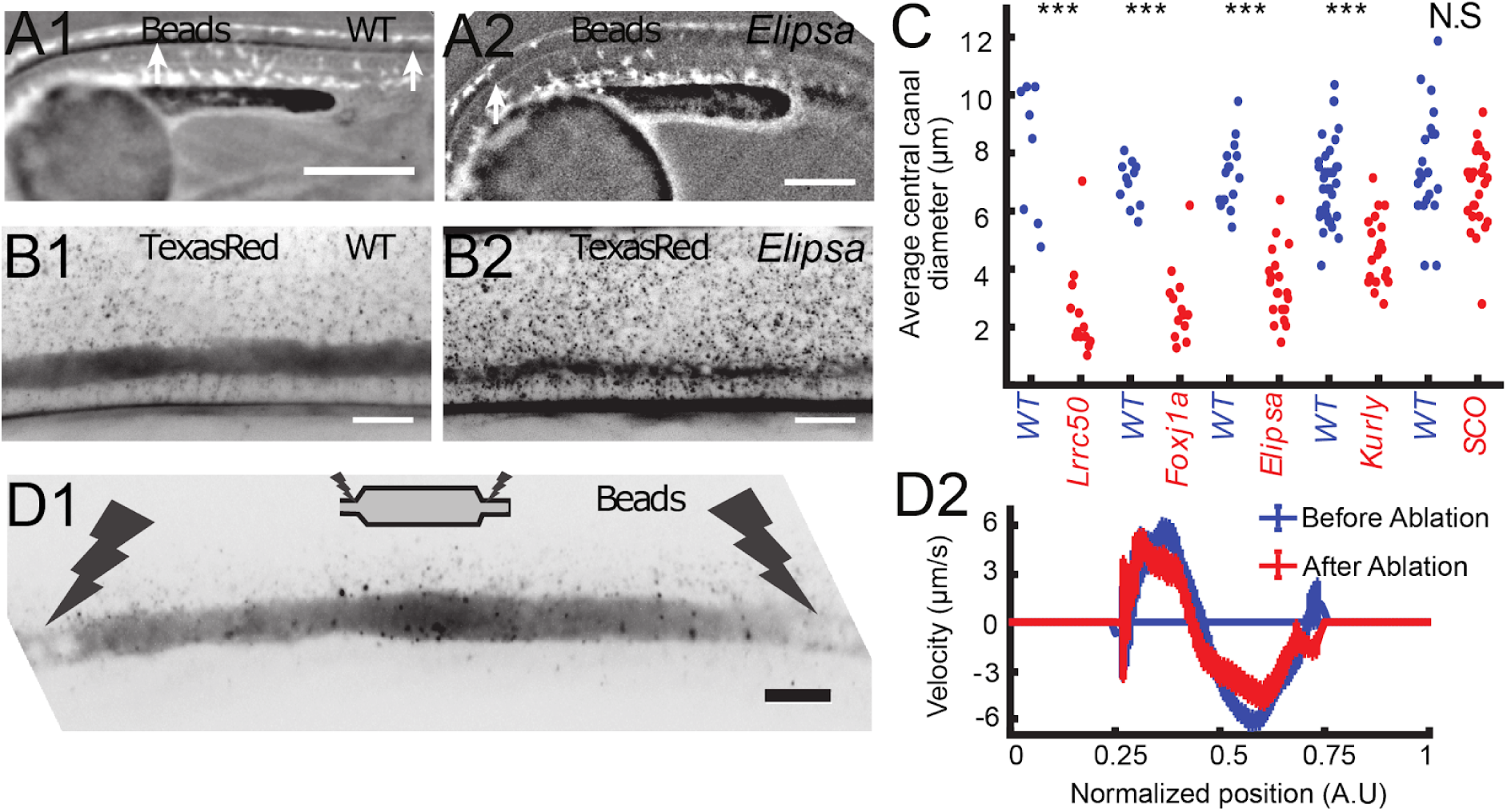
Cilia motility controls long-range transport, central canal architecture, and local flow. (A1, A2) Progression of 20 nm fluorescent beads inside central canal 2 hours post injection (2 hpi) in the diencephalic ventricle in a WT embryo (A1) and in a ciliary mutant (elipsa) sibling (A2). The progression front is emphasized by the rightmost white arrow in (A1) and (A2), showing that beads little propagate in central canal in ciliary mutants. (B1, B2) Central canal geometry in a WT embryo (B1) and in a ciliary mutant (*elipsa*) sibling (B2). The central canal was filled with Texas Red, and displayed as the maximal projection image from a Z-stack of 30 images around central canal. The latter appears collapsed in an *elipsa* mutant. (C) Quantification of central canal diameter for different ciliary mutants (*lrrc50, foxj1a, elipsa, kurly*) and one curled down mutant with intact CSF flow (*SCO*) (red dots, right side) versus their WT siblings (blue dots, left side). (D1) Time average image of 20 nm fluorescence beads in inverted contrast of a central canal region where photoablation was performed on both sides (dark lightning), closing the canal. (D2) Quantification of local CSF flow in the central canal region in (D1) before (blue) and after (red) photoablation. Central canal geometry remains intact in the central region (D1), as well as the flow profile (D2). Scale bars represent 200 μm in (A1, A2), and 15 μm in panels (B1, B2, D1). P-values in (C) are respectively 2.2 10^-6^, 3.1 10^-9^, 3.5 10^-6^ 1.82 10^-8^, 0.096 from left to right. The values for control WT siblings from all experiments are not statistically different.

Because exogenous beads are not transported down the CC in paralysed mutants, one cannot directly test the role of motile cilia on generation of the local CSF flow. Instead, we investigate whether letting mutant embryos with ciliary defects spontaneously twitch would facilitate the transport of beads. Indeed, we find that for mutants with a moderate reduction of the CC diameter, such as *elipsa* and *kurly*, some beads can enter the CC. These beads exhibit Brownian motion and some exhibit a residual flow with numerous recirculation zones (**Supplementary Movie S4**), in good agreement with our “scarce cilia” model.

Our model also predicts that the bidirectional CSF flow in the CC is generated locally and does not rely on a global pressure gradient generated in ventricles. This is confirmed by multiple observations: *i)* even after large regions without cilia and local flow, the CSF flow in more caudal regions can show a bidirectional flow, as long as motile cilia beat locally in the ventral region (see **Figure 3c**). *ii)* When some portion of around 200 μm of the CC is isolated by using photoablation to block flow on both sides (**Figure 4d1**), the canal remains wide-open and the CSF still flows bidirectionally (**Supplementary Movie S5**) with a similar velocity profile (**Figure 4d2**).

### Muscle contraction transiently modifies CSF flow

So far, we mainly restricted our study to paralyzed zebrafish embryos. However, muscle contractions are also at the origin of several biological flows^47^, and were recently shown to create a massive transient flow in the brain ventricles of zebrafish embryos^20^. Consequently, we investigate in details the CSF flow and transport on spontaneously twitching embryos restrained in agar where contraction can be identified from the notochord displacement (white arrow) on the transmitted image (**Figure 5a**). Following each contraction, rapid displacements of CSF (too fast to be properly captured here) are observed in either caudal or rostral direction, followed by a slower counter-flow in the opposite direction, whose velocity decays until the original bidirectional flow is recovered (**Figure 5b, 5c**, **Supplementary Movie S6**). The average CSF flow through the entire CC section is correlated with the instantaneous muscle contractions (**Figure 5d**) showing a transient increase after contractions. By comparing the increase in total CSF flow over 5 seconds after each contraction versus the contraction strength, a robust increase in flow rate occurs after each contraction (**Figure 5e**; 204 contractions from 22 WT embryos). However, we do not find any obvious correlation between the flow direction and our estimates of the contraction direction or strength. Body contractions are also shown to enhance transport of beads in mutants where cilia are defective (**Supplementary Movie S4**). As no significant difference in the CC diameter is detected during the muscle contractions, we propose that the muscle-dependent enhancement of CSF flow is not due to a local pinching of the canal, but rather to a change in brain ventricles volume recently observed^20^.

**Figure 5.**
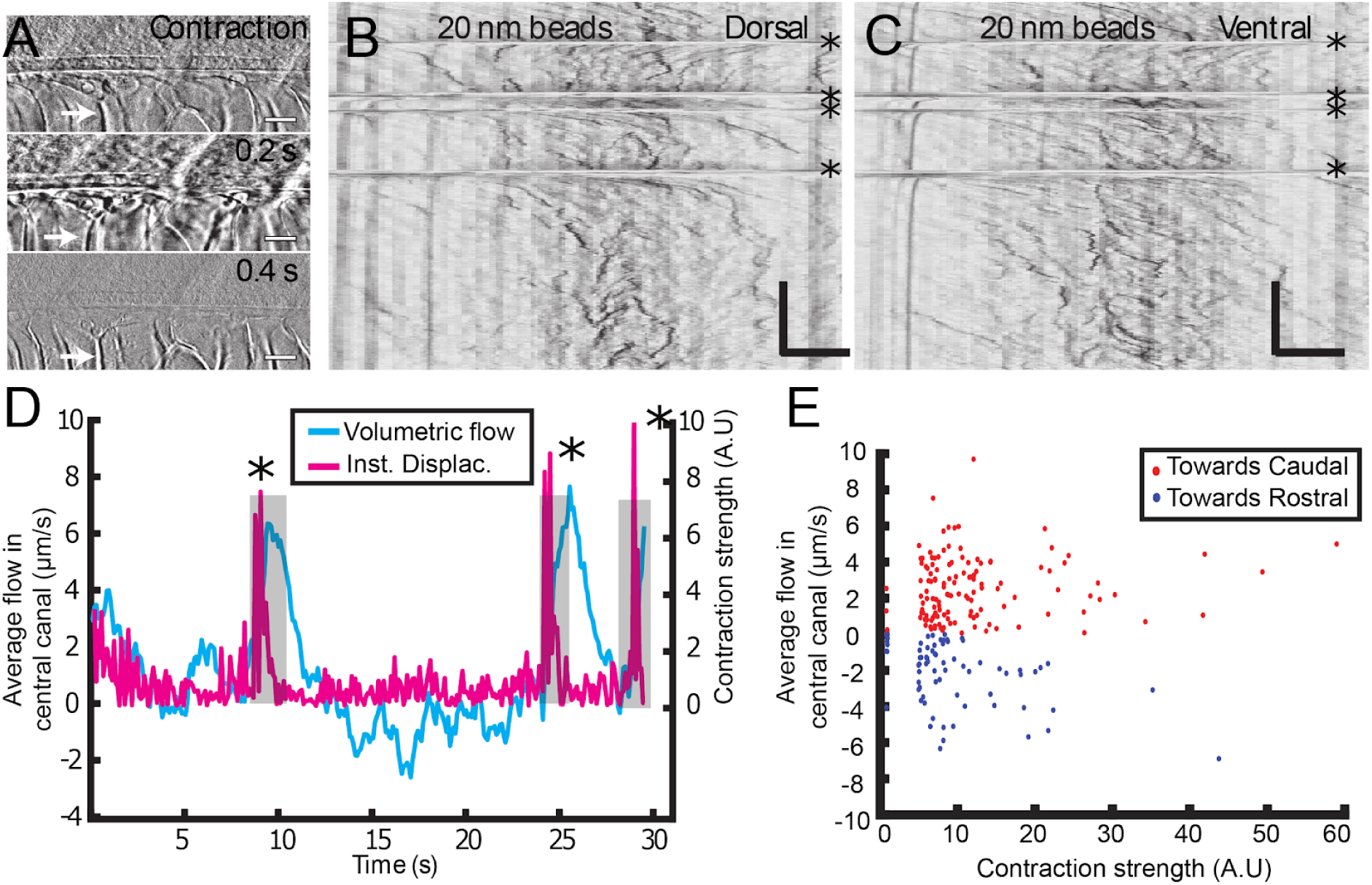
Muscle contraction transiently increases local CSF flow. (A) 3 successive images showing the time derivative of the transmitted intensity of a zebrafish embryo during muscle contraction. A fast contraction in one direction is followed by a slower decay back to the initial position, during which the notochord comes back slower to its original position (white arrows). (B, C) Dorsal (B) and ventral (C) kymographs obtained after spinning disk imaging of injected 20 nm beads. Contractions can be identified as the sharp horizontal lines and are emphasized by the four black asterisks. (D) Average volumetric flow across one section of the central canal versus time (cyan curve) for one representative embryo. The embryo instantaneous displacement (magenta curve) is superimposed. Contractions are also emphasized by dark asterisks and a grey background. (E) Quantification of the sum (integral) of the average volumetric flow during 2.5 seconds after each of the 204 contractions from 22 WT embryos. These values are plotted versus the contraction strength, *i.e.* the integral of intensity derivative during the time of contraction. The dataset is separated in positive flows (red dots), where flow after contraction is directed towards the tail, and in negative flows (blue dots), where it is directed towards the head. Horizontal scale bar is 15 μm in (A, B, C). The vertical bar in (B, C) is 4 seconds.

### Bidirectional long-range transport in the central canal is accelerated by the flow

Now that the CSF flow has been fully characterized, its ability to transport particles on long distances needs to be demonstrated. The long-range transport follows an apparent diffusive law, as the propagation front of nanoparticles progresses with the square root of time^36^ and depends on particle size (**Figure 6a1, 6a2**). The propagation front is also accelerated by muscle contractions (**Figure 6a3**). In practice, 20-nm particles injected in the DV travel through the entire CC in around two hours (**Figure 6a2**), far from the timescales expected for both passive diffusion (more than one day) and sole active transport of 5 μm/s (less than 10 minutes). This suggests that the transport of particles should be modelled by including an advection term in the diffusion equation. In a bi-dimensional canal (**Figure 3a2**), the transport equation can be written as:

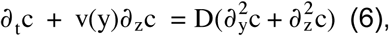

where c is the local concentration of particles, and ∂_t_, ∂_z_ and ∂_y_ are respectively the time and spatial derivatives in the rostro-caudal (z) and dorso-ventral (y) directions. v(y) is the velocity profile (**eqs. 1, 2**), and D is the diffusion coefficient of particles in CSF. For a spherical particle of radius r, the Stokes-Einstein relation gives:

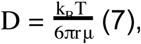

where T is the temperature and k_B_ is Boltzmann’s constant.

**Figure 6.**
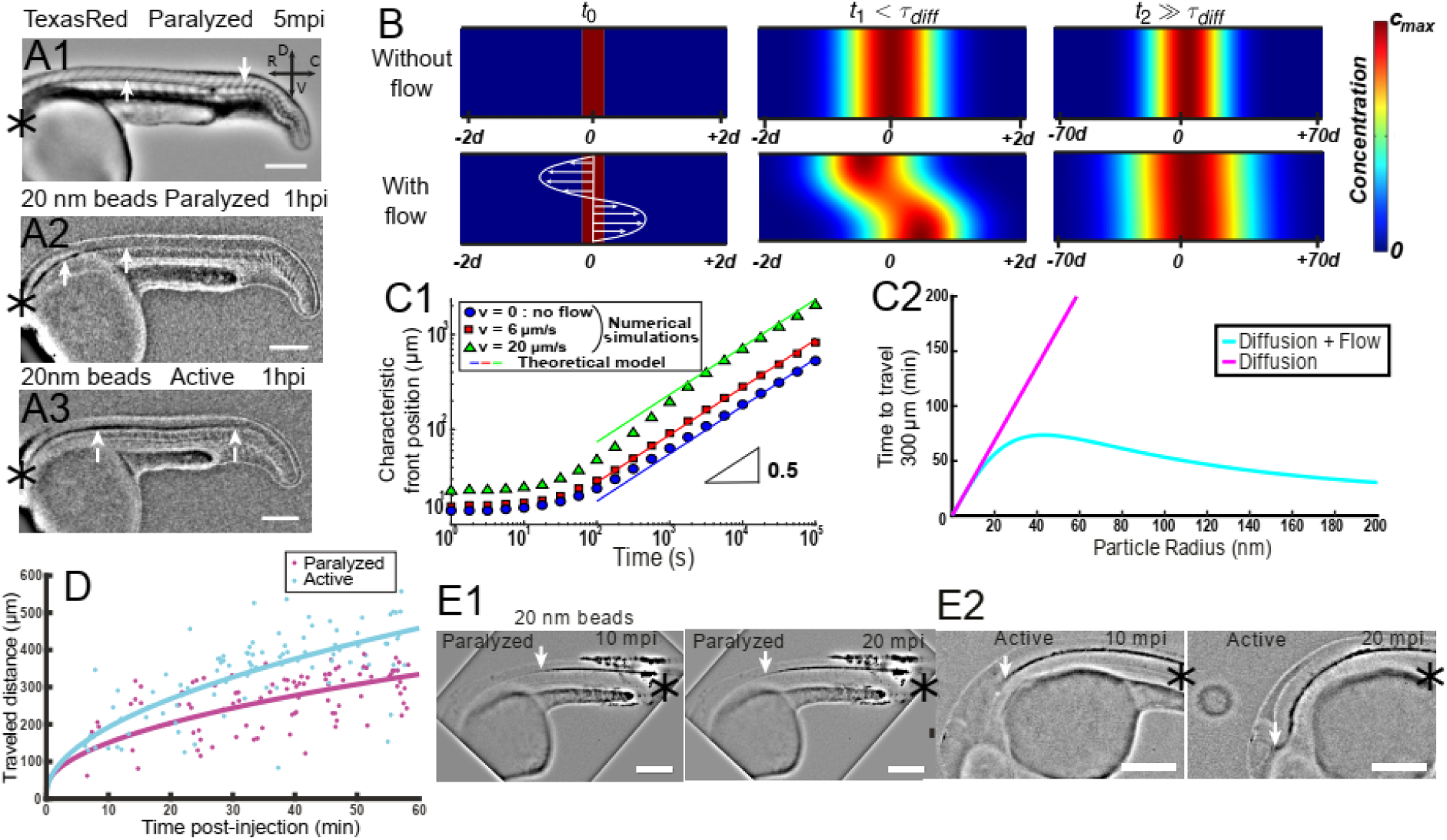
Local CSF flow accelerates the bidirectional long-range transport along rostro-caudal axis. (A1) Inverted contrast fluorescence widefield image of the propagation front of Texas Red 5 minutes post-injection (mpi). (A2, A3) Inverted contrast fluorescence widefield image of the propagation front of 20 nm beads 1 hour post-injection in the DV of a paralyzed (A2) and active (A3) embryo. White arrows in (A1, A2, A3) show central canal and the propagation front. (B) Numerical simulation of Taylor Aris diffusion, accounting for long-range transport in central canal. A small region of high concentration is created at the beginning of the simulation (t_0_ – left column) with (bottom) and without (top) the flow. The propagation front first follows the local flow profile (t1, center column), but a Gaussian distribution is finally recovered with and without flow (t2, right column). However, the concentration front has propagated faster in the presence of flow. The color represents the local concentration in particles. (C1) Numerical results of the characteristic front position of 20 nm radius particles versus time for different bidirectional flow velocities in log scale (v=0, 6, and 20 μm.s^-1^ for respectively the blue dots, the red boxes, and the green triangles). The colored solid lines correspond to the asymptotic prediction (0.5 line) of pure diffusion. (C2) Theoretical calculation of the average propagation time of a solution of spherical particles to travel 300 μm in central canal versus the particle radius with (cyan) and without (magenta) the bidirectional flow. (D) Experimental propagation front of injected 20 nm beads versus post-injection time, measured in 16 paralyzed (magenta) and 16 active unparalyzed (cyan) embryos. (E1, E2) Inverted contrast fluorescence widefield image of the propagation front of 20 nm beads injected in the caudal central canal of a paralyzed (E1) and active zebrafish embryo (E2), showing “reverse” (caudal to rostral) transport between left (10 mpi) and right images (20 mpi). The black asterisks in (A1-A3, E1, E2) show the injection site.Scale bar is 200 μm in (A1, A2, A3, E1, E2).

As seminally described by Taylor^48^ and Aris^49^, the diffusivity of solutes is amplified in presence of velocity gradients, due to a coupling between pure diffusion and flow-induced transport. They demonstrated that on long distances, the transport still follows a diffusive process, with an effective diffusivity enhanced by the shear flow. Following analogous developments of Bruus^50^, we perform the theoretical derivation of the effective diffusivity D_eff_ of particles submitted to a bidirectional flow (**Supplementary Information**), which leads to:

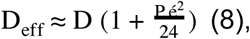

where P é = Vd/D is the dimensionless Péclet number, comparing the characteristic times for spanwise diffusion d^2^/D and convection d /V. This theoretical prediction is validated with numerical simulations, comparing the long-range transport of particles of diameter 40 nm (in the range of commonly observed embryonic extracellular secreted vesicles in CSF^29^), without flow, and in presence of a bidirectional flow (**Figure 6b**). The simulations show that after a time larger than the characteristic diffusion time d^2^/D= 10 s, a diffusive-like gaussian concentration profile is recovered, with a wider extent in presence of a flow (**Figure 6b**). This diffusive transport is numerically obtained for three different maximal speeds of the flow: 0, 6 μm/s, and 20 μm/s (**Figure 6c1**), and shows that transport is accelerated by the flow speed, as predicted by our theoretical derivation. Another interesting feature of this diffusivity enhancement is its weak dependence on the size of the transported particles, as opposed to classical diffusion for which the diffusivity is inversely proportional to the particle size (**Figure 6c2, eq. 7**).

*In vivo* front tracking experiments illustrate this flow-induced accelerated transport of 20 nm nanoparticles in paralyzed and active zebrafish embryos (**Figure 6d**). While both active and paralyzed embryos exhibit a diffusive-like transport, the effective diffusivity is higher in the case of contracting embryos. It confirms that muscle contractions can increase the long-range transport efficiency, which act similarly to a constant increase of the local flow velocity (**Figure 6c1**). Interestingly, our model also predicts that transport is enhanced in both rostro-caudal and caudo-rostral directions. We indeed demonstrate that beads injected locally in the caudal part of the central canal are transported rostrally towards the brain ventricles (**Figure 6e1**). Similarly to rostral-to-caudal transport, the reverse transport is sped up for contracting embryos (**Figure 6e2**).

### The connection from brain ventricles to central canal in the spinal cord is gated by a funnel-shaped channel and is critical for embryonic growth

Now that the ability of the CSF flow to transport particles across the whole CSF zebrafish body has been demonstrated, its consequences for embryonic development have to be discussed. We first summarize the entire CSF circulatory system of zebrafish embryos (**Figure 7a**). The DV is connected to the CC via a small channel making a funnel connection of small constant diameter (0.55 ± 0.12 times the CC diameter and 233 ± 24 μm long, **Figure 7b1, b2, b3**, 11 embryos). At the caudal end of the CC, a secondary channel thinner than the CC closes the loop by connecting the CC to the rhombencephalic ventricle (RV) (**Figure 7a**). In 24 hpf to 30 hpf embryos, primary cilia are found in the brain ventricles but they are mostly non-motile. Therefore brain CSF flow is mostly brownian with a pulsatile component at the heart beat frequency, which is hindered if the heart beat is suppressed (**Supplementary Movie S7**), indicating that brain ventricles marginally participate to the global circulation of CSF between 24 and 30 hpf. On the contrary, a strong unidirectional flow towards the caudal region is driven through the funnel (**Supplementary Movie S8**), with a few smaller channels connecting the funnel to the bottom of the RV. Through the funnel, this fast unidirectional flow is characterized by parabolic Poiseuille-like velocity profile of maximal speed 15 μm/s (**Figure 7c1, 7c2**). We observe motile cilia inside the funnel (data not shown), but the constrained geometry prevents us to analyze cilia distribution and properties in this channel.

**Figure 7.**
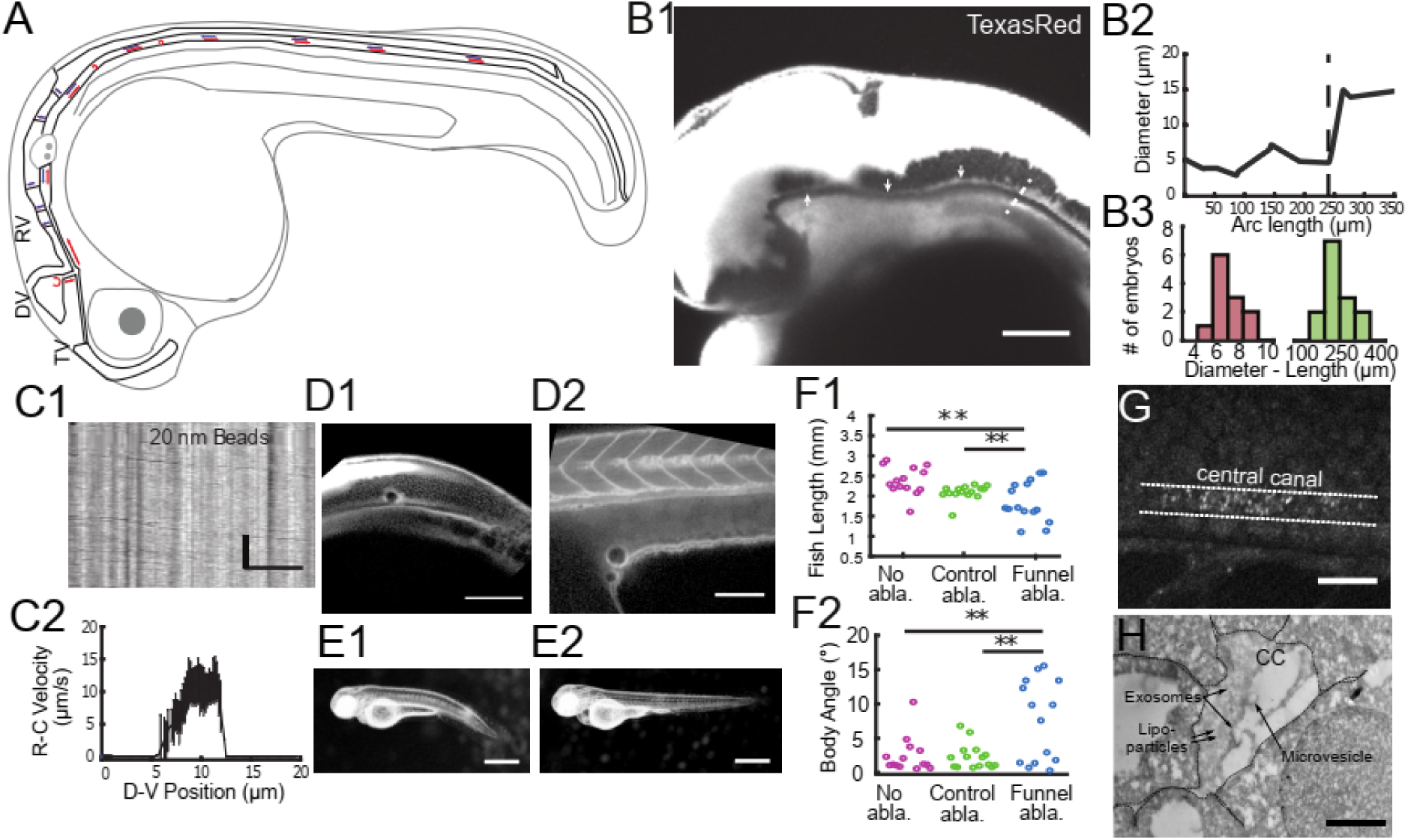
Funnel between brain and spinal cord controls growth during embryogenesis. (A) Scheme of CSF circulation in 30 hpf zebrafish embryos with CSF-filled cavities emphasized by black lines. Red arrows represent net flow towards the tail, while blue arrows represent net flow towards the ventricles. Red turning arrows illustrate the presence of vortices. (B1) Confocal image of ventricles and funnel after Texas Red injection. The signal from the ventricles is highly saturated to enable visualization of the funnel (white arrows, before white dashed line). (B2) Experimental measurement of the funnel diameter (in (B1)) versus its arc length. The diameter shows a sharp increase at the end of the funnel (black dashed line). (B3) Histogram of the average funnel diameter and funnel length measured in 11 WT embryos, giving respectively 7.1 ± 0.7 μm (ratio with central canal diameter of 0.55 ± 0.12, as discussed in Methods) and 233 ± 24 μm (median ±0.5 s.d.). (C1) Inverse contrast kymograph of 20 nm beads flowing in the funnel after injection in the diencephalic ventricle. (C2) CSF flow profile inside the funnel, captured with our automatic kymograph analysis. (D1, D2) 2-photons image of fluid-filled cavities after Texas Red injection and after optical ablation at the end of the funnel (D1), and in the yolk extension (D2 -control ablation). (E1, E2) Widefield transmitted images of 3 dpf zebrafish larvae 30 hours after funnel optical ablation (E1), or control ablation (E2). (F1, F2) Quantification of the fish total length (F1), and the main body angle (F2), 30 hours after Texas Red injection without ablation (magenta), after control (green) and funnel (blue) ablation with 14 larvae per category. P-values for length and angle are respectively 1.3 10^-3^ and 3.0 10^-3^ between funnel and no ablation, 9.6 10^-3^ and 6.4 10^-3^ between funnel and control ablation, and 0.066 and 0.89 (not significant) between the two controls. (G) Full field optical coherence tomography imaging of 30 hpf embryos reveals high density of secreted vesicles in central canal. (H) Electron microscopy with immunogold labeling of extracellular vesicles (EVs) in central canal of a 1 dpf zebrafish larva. Scale bar is 100 μm in (B1, D1, D2), 15 μm in (C1, G), and 500 nm in (H). The vertical scale bar in (C1) is3 seconds.

The funnel allows a connection between the DV and the CC during development, that we investigate by performing 2-photon mediated photo-ablations of the connection between the funnel and the CC (**Figure 7d1**), and control ablations in the yolk extension (**Figure 7d2**). 30 hours after ablation, half of funnel-ablated larvae were shorter than siblings undergoing control ablation or without ablation (**Figure 7e1, 7e2, 7f1**) and they often exhibited an increase in body-axis curvature (**Figure 7f2**). We thus investigate whether the CSF contained extracellular vesicles, using label-free imaging technique, called full field optical coherence tomography (**Figure 7g** and **Supplementary Movie S9**). We identify many vesicles with high lipids or proteins content floating in central canal, in agreement with previous observations^31^. The presence of dense exosomes is confirmed using fluorescence labeling (**Supplementary Movie S10, Methods**), and electron microscopy with immunogold labeling of exosomes (**Figure 7h, Methods**)^30^. Note that both optical techniques show similar bidirectional flow of endogenous extracellular vesicles in the CC (**Supplementary Movies S10, S11**). Altogether, these experiments indicate that CSF circulation contributes to embryonic growth, and suggest that the transport of extracellular vesicles between brain and spinal cord is a critical process.

## Discussion

In this study, we have performed a novel quantification of cerebrospinal fluid (CSF) circulation in zebrafish embryos between 24 and 30 hpf, demonstrating for the first time that a constant bidirectional CSF flow is driven by ventral motile cilia and that body movements create additional transient flows. These two driving forces accelerate the long range transport of all particles larger than 10 nm, including macromolecules and extracellular vesicles present in the CC. This permits fast transport both from brain to spinal cord, and from spinal cord to the brain, apparently critical for embryonic development. In addition, we reported two intriguing phenomena: 1) in mutants with ciliary defects, the CC collapses a phenomenon that had been missed in previous characterizations; 2) the density of extracellular lipidic vesicles in CC, possibly containing exosomes, is particularly high at the embryonic stage, as already reported in mammals using fixed tissue or CSF punctures^27,29^. We showed using 2 live imaging techniques that extracellular lipidic vesicles are transported in both directions along the rostro-caudal axis.

Little is known about CSF embryonic flow as its investigation represents a daunting imaging challenge. Indeed, CSF is a clear fluid with little endogenous contrast, and the CC is a small channel difficult to access inside the spinal cord. In healthy young humans and mammals where the CC is open, the canal is often too small to be imaged with phase-contrast MRI^51^, in contrast to brain ventricles or subarachnoid spaces. Here, we take advantage of the transparency of zebrafish embryos to characterize CSF bidirectional flow as well as cilia dynamics in the CC confined geometry. A simple hydrodynamic model accounts for our observations by modeling cilia contribution as a homogeneous volume force f_v._ The latter fits experimental data for an amplitude of 4000 N/m^3^, corresponding to a force of 10 pN generated by a single cilium occupying a volume of characteristic size 10 μm, in good agreement with the values reported in the literature^52^. This modeling of the force f_v_ is reminiscent to the study of Siyahhan *et al.*^*^53^*^ in ventricles of the human brain, where a linearly varying force of comparable magnitude is used instead. We report that including a linear force profile does not significantly alter the predicted velocity profile (**Supplementary Figure S2b**), which shows the robustness of these averaged models. We acknowledge here that we cannot model the complexity of many interacting cilia beating at many different frequencies, and that only the average flow can be inferred by our model. Interestingly, our model can efficiently predict flow profiles for many other aspect ratios between the cilia extension and the channel diameter and heterogenous ciliary beating frequency (**Supplementary Figure S2c**). Notably, it convincingly predicts the flow previously reported by Shields *et al.* in the context of artificial magnetic cilia in microchannels ^8^, where this aspect ratio is low. Our model should also be relevant to account for cilia-induced flows in brain ventricles or in their connecting channels where the confinement is moderate.

Our model also predicts the establishment of an overpressure in the CC, increasing linearly towards the caudal direction (**Supplementary Figure S2a**). This inner pressure is dramatically decreased as the density of motile cilia is lower. These findings may be directly related to the collapse of the CC reported for mutants with defective cilia (**Figure 4c**). This suggests that, among other roles related to transport of chemical cues or mechanosensing, the cilia-induced flow ensures the integrity of the CC, necessary for a proper body axis formation. Interestingly, similar transitions from cylindrical to collapsed geometries due to inner pressure decrease were already reported in small veins^1^. It may also account for the apparent fragility of the CC that easily collapses during fixation (Wyart lab, unpublished observations). We report that mutants with defective cilia were unable to maintain an open CC. *Elipsa* and *kurly* embryos with partial loss of cilia motility show intermediate phenotypes, with some regions open and other collapsed, and with open regions showing bidirectional flow with a higher density of recirculation zones. We believe this intermediate phenotype is predicted by our “scarce cilia” model (**Figure 3c**), since a few cilia remain motile in *kurly* embryos^54^ and in *elipsa* between 24 hpf and 30 hpf^39^. The model also accounts for the reduced long-range transport previously observed in *kif3b* mutants, where cilia density is reduced^36^. In contrast, *foxj1a* mutant embryos showing ependymal cells with defective cilia^55^ (**Supplementary Figure S3**) have a completely collapsed CC. This observation strongly suggests that the motile cilia generating the flow and maintaining the CC geometry originate from ependymal cells only, while remaining cilia probably belonging to CSF contacting neurons^56^ are too short and not dense enough to generate a stable CC (**Supplementary Figure S3**).

We provided the first description of the narrow channel linking the CC with brain ventricles with a funnel-shaped entrance from the diencephalic ventricle. We found another channel connecting the caudal part of the CC to the rhombencephalic ventricle, forming a closed loop (**Figure 7a**). The pressure distribution in the CC induced by the motile cilia could thus also be responsible for a flow in these narrow channels even if they would not contain any motile cilia. The presence of motile cilia, observed in the funnel, may accelerate the flow even further. The cilia in the CC have consequently the potential to drive a large scale pressure-driven flow as well, playing the role of a pump for the whole CSF circulatory system. Unfortunately, flow measurements in these narrow channels are extremely challenging and will be the scope of a future study. Based on our ablation experiments we propose here a critical role for the passage of CSF from the brain to the spinal cord through the funnel, during embryonic development. This correlates well with previous reports of body axis straightening and spine morphogenesis defects arising when the CSF flow in ventricles is altered at later stages^21,36,37^. Another interesting follow-up, today limited by our imaging technology, is to investigate the role of body movements in freely behaving embryos, knowing that they normally twitch every few seconds.

In this study, we neglected the presence of the Reissner’s fiber (RF) in the central canal, although it was shown to play a critical role in body axis straightening^39^. As it does not compartmentalize the fluid in 3D, the presence of the RF does not qualitatively modify the nature of the flow and its bidirectionality, as confirmed by velocity profiles measured in *scospondin* mutants deprived of RF^39^. However, we expect an increase in the viscous stress at the vicinity of the fiber thereby likely changing the mechanical force applied by CSF flow on cells contacting the CC. As CSF-contacting neurons are known to be mechanosensitive^31,57^, a change in activity of these neurons in the absence of the RF may also occur.

Finally, our study focused on embryogenesis as this is a critical period when CSF shows its highest concentration in secreted extracellular lipidic vesicles^27,29^, and when blood flow along the body and ciliary beating in the ventricles are not yet implemented. As the structure of the central canal is well conserved among vertebrates^58^ but very challenging to access with optical imaging in most species, we expect that our findings in zebrafish will guide future investigations in mammals. The thorough study of CSF flow and role in larval, juvenile and adult stages will be the object of future developments, as we expect the CSF flow to be different at later stages since both structure and cilia dynamics in the CC change with time. Altogether, we believe our work to be a major first step into the understanding of the role of CSF transport in central canal to ensure a long-range communication between cells throughout the body that may control development, inflammatory response, and modulation of locomotor and postural functions carried by spinal circuits.

## Materials and Methods

### Table with fish mutants and transgenic lines used in this study

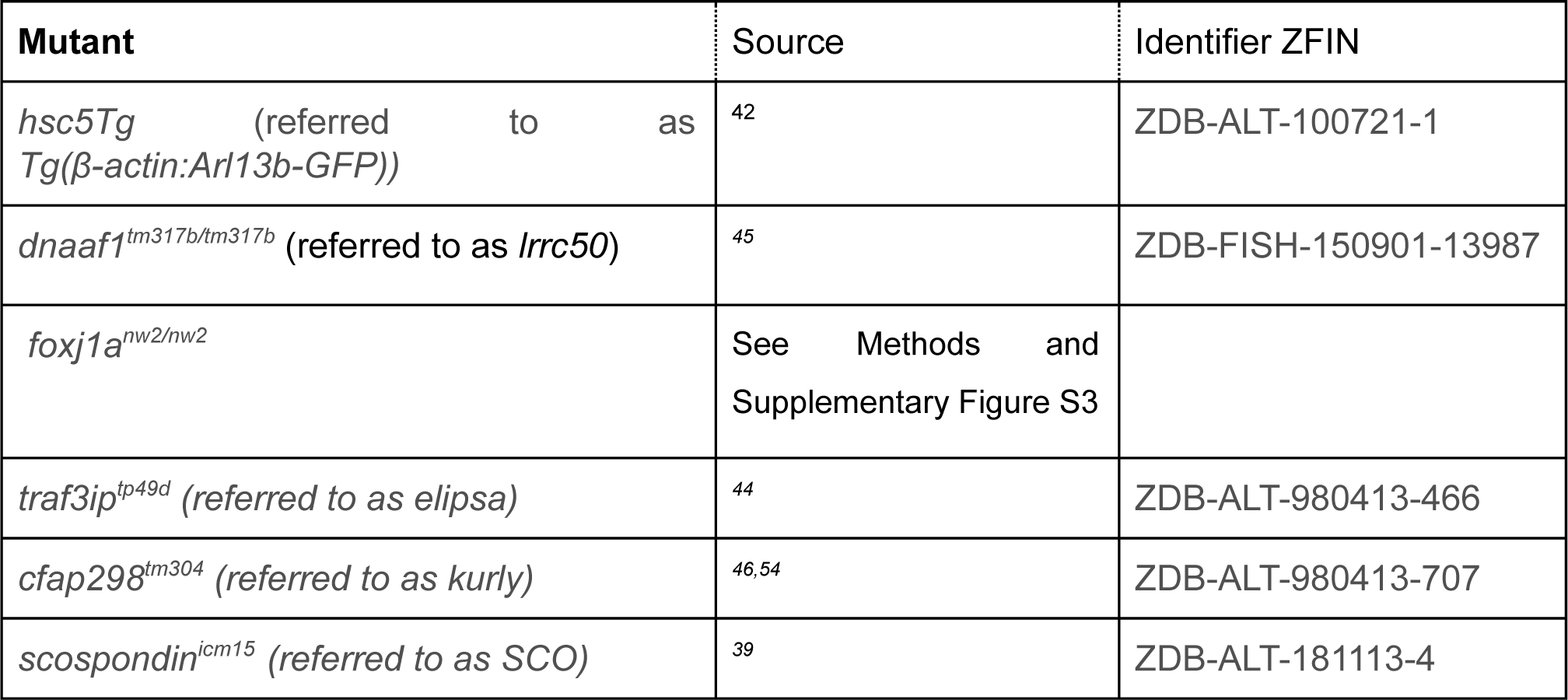

### Animal husbandry

All procedures were performed on zebrafish embryos before 2 days post fertilization in accordance with the European Communities Council Directive (2010/63/EU) and French law (87/848) and approved by the Brain and Spine Institute (Institut du Cerveau et de la Moelle épinière). All experiments were performed on *Danio rerio* embryos of the AB strain, and nacre background. Animals were raised at 28.5°C under a 14/10 light/dark cycle until the start of the experiment.

### Quantification and Statistical Analysis

All values are represented as median ± 0.5 standard deviation (s.d.) to evaluate the spread of each distribution, except for the error bars in the flow profiles showing the standard error of the mean (SEM), in which case we are interested in the average flow at one position. Hence, the speed distribution of particles is not per se interesting as it depends on experimental errors and Brownian motion. All statistics were performed using MATLAB. Statistical details related to sample size, and p values, are reported in the figure legends. In figure panels, asterisks denote the statistical significance calculated by two-tailed t test for samples with unequal variance (Welch t test): ^*^, p < 0.05; ^**^, p < 0.01; ^***^, p < 0.001; ns, p > 0.05.

### Injection in the CSF

30 hpf embryos were manually dechorionated, mounted in 1.5% low-melting point agarose in lateral position. A few nanoliters of a mix containing 20 nm beads, 3,000 MW dextran, labeled with Texas Red, and alpha-bungarotoxin (TOCRIS) were injected in the center of the diencephalic ventricle in order to label CSF and paralyze the fish in a single injection. The injections were performed using a Picospitzer device (World Precision Instruments). 20 nm carboxylate FluoSpheres of center wavelength 505/515 nm (yellow/green, F8888, Molecular Probes©) were diluted to a 2% concentration in artificial CSF (containing in mM: 134 NaCl, 2.9 KCl, 1.2 MgCl2, 10 HEPES, 10 glucose, 2 CaCl2; 290 mOsM +/- 3 mOsm, pH adjusted to 7.8 with NaOH), and then sonicated for 10 seconds. 3,000 MW dextran labeled with Texas Red (Ex/Em wavelength: 595/615/ ThermoFisher Scientific D3329) was diluted at 0.2 % mass/volume concentration and was used both to serve as a marker of injection, and to assess the boundaries of the ventricles and central canal. A final concentration of 100 μM of alpha-bungarotoxin was used. We could visually assess that such concentration, as well as the ventricle injection of alpha-bungarotoxin, although not conventional, were still efficient enough to paralyze the embryos. For experiments in contracting fish, the alpha-Bungarotoxin was replaced by artificial CSF. Before performing CSF flow measurement in central canal, a delay of one hour was taken in order for the particles to be transported down the central canal. Beads larger than 100 nm-diameter beads were restrained to brain ventricles, while we could beads of 20 and 45 nm diameter entering the central canal.

### Fluorescent beads imaging

For CSF flow experiments, time-lapse images were acquired at room temperature on an inverted Leica DMI8 spinning disk confocal microscope equipped with a Hamamatsu Orca Flash 4.0 camera, using a 40X water immersion objective (N.A. = 0.8, pixel size 189 nm). 300 images of the beads mostly in the rostral part of central canal were acquired at a frame rate of 10 Hz for 30 seconds using Metamorph software (www.moleculardevices.com). Some experiments in contracting fish were performed at a frame rate of 40 Hz for 20 seconds to be able to follow more accurately the embryo contraction.

### Particle Tracking velocimetry

A custom Matlab Particle Tracking Velocimetry (PTV) algorithm was adapted from a classical PTV algorithm^59^ and used to obtain instantaneous particle velocities. This algorithm was only used to produce Fig 1b as it only worked consistently in the best SNR conditions. The first crucial step was to segment the particles inside the region of interest corresponding to the central canal. This region was automatically inferred based on an intensity threshold on the high pass filtered mean image. Particles segmentation was more efficiently performed using a machine learning based segmentation software (iLastik^60^). Particle fine positioning was performed by fitting a 2D gaussian to each segmented region (particle) and by extracting the center position. A large matrix of particle position (X, Y, time) is obtained at this point. Then, the PTV algorithm itself is applied. It consists in connecting the particles found at successive times in order to minimize the total squared distance traveled by all beads. Note that this algorithm is more optimized to purely Brownian displacements and limit the number of particles that one can track before saturating its computer memory, as the number of calculation scales as N_particles_^2^. Once the particles have been tracked, each trace is obtained and kept only if it is longer than 5 time points. The instantaneous speed is calculated and plotted as a vector on the mean intensity image. An additional color code ranging from blue to red corresponding from the minimal to maximal speed.

### Automatic kymograph measurement of beads velocity and CSF flow profile

First, time lapses were automatically rotated and cropped with an extra manual step for fine adjustments to isolate a portion of the central canal and orient it in a correct dorso-ventral orientation. A 2D wavelet filter (decomposition of level 6 with haar wavelet coupled to a 2-D denoising scheme based on level-dependent thresholds penalyzing low signal pixels) is applied to increase the local signal-to-noise ratio of the fluorescent beads. Stacked images were then re-sliced and particle trajectories were automatically tracked and analyzed for every successive dorso-ventral positions. The average signal on 3 consecutive positions is calculated with a moving-window average. Each column is then divided by its average value to prevent from spatially or temporally inhomogenous light illumination and photobleaching if any. Only pixels above the average signal are kept, and an internal Matlab segmentation algorithm (regionprops) is applied to extract regions of interest in each kymograph. In order to filter only the straight lines corresponding to particle motion, a few additional empirical filtering conditions were applied: Only the regions with enough pixels (15 pixels at least to avoid random fluctuations), corresponding to lines (eccentricity above 0.9 to prevent from selecting big aggregates or any large objects) were kept. Additionally, to prevent selecting particles stuck in the canal (vertical line in the kymograph), or camera/illumination fluctuations (global change of intensity in the entire image, leading to an horizontal line in the kymograph), we filtered any region whose orientation is close to 0 or 90° ± 180° (excluding regions whose cosine or sine of orientation are below 0.1). Note that the intensity threshold is not so strong, but that most of the particle filtering is made through these 4 last conditions. A few regions corresponding to individual particle rostro-caudal trajectory are thus obtained. The tangent of these lines orientation is calculated, and once corrected by pixel size and frame time, the individual particle velocity is obtained. All particles from one kymograph were aggregated to the average velocity and standard error to infer the local flow (which we are interested in rather than individual particles speeds). Note that we refer to the number of events, or number of lines tracked, rather than the number of particles, as this algorithm cannot guarantee that one particle is not tracked several times if its dorsoventral position changes over time.

### Extraction of parameters in the velocity profiles

Once the profiles are obtained, we extract some parameters in order to compare the profiles from many embryos. The maximal and minimal values, as well as their respective positions were extracted. The position at the center of the profile where the speed shift from a positive to a negative value is also extracted. The position in the canal is normalized between 0 and 1, where 0 is the most ventral position where at least 5 moving particles were found and velocity exceeds 0.4, and 1 the most dorsal position where 5 moving particles were found and the velocity is not higher than −0.4.

### Flow versus time analysis

In Figure 5d, the average flow versus time is plotted in embryos able to contract in agarose. Protocol and analysis are similar than for the flow profile analysis, except that alpha-bungarotoxin was removed from the injection mix and that the dataset is analyzed in a different manner. The same automatic kymograph approach is used, in order to obtain a number of events corresponding to beads motion. For each event, each trace is kept and associated to its speed, its central position in D-V and R-C axis, as well as its time of occurence. If the speed profile is obtained by averaging over time for all pixels along the D-V axis, the average flow versus time is obtained by averaging all particles from all D-V positions at a given time. When bidirectional flow is achieved (before contraction, or in a paralyzed embryo), the average flow is close to 0. We could in theory perform as many refinements as required, e.g. averaging on only all dorsal or all ventral particles instead of all particles (not shown here). The only constraint is to have enough events at each timepoint, so that the flow measurement is meaningful and the Brownian part of particle motion is averaged out. Also in Figure 5d, the fish contraction strength is estimated to be correlated to the temporary increase in volumetric flow. To estimate such contraction, the average signal from all pixels within central is calculated and the absolute value of their time derivative calculated (magenta curve in Figure 5d). An arbitrary threshold of 4 was empirically chosen to select points during contraction. The contraction strength used in Figure 5e is calculated as the sum of the displacement signal during one event with displacements values above threshold. This value voluntary mix the instantaneous derivative value and the contraction duration, as they can both play a role in the establishment of a different flow. In datasets where embryos did not move, the average flow and displacement were calculated over the entire time serie. The average displacement value was calculated in order to avoid getting a large value due to the large time span of the dataset (and due to the use of absolute value).

### Comparison between methods to measure local flow (PTV/PIV/and kymograph approach)

Besides the PTV and kymograph approaches, we also tried to use a particle imaging velocimetry (PIV) algorithm, which is generally used to measure local flow^61,62^. This section aims at comparing all these techniques qualitatively with respect to our imaging conditions. Before describing why we mainly used the automatic kymograph approach, we have to recall that we cannot inject the particles directly in the central canal, and realized that exogenous particles above 100 nm in diameter stayed confined inside the brain ventricles. In turn, we could only image particles of diameter below 100 nm, and mainly used 20 nm particles with fairly low signal, especially at high imaging frequencies. Besides, the Brownian motion of 20 nm beads is important, and can generate instantaneous displacements of the same order of magnitude than the CSF flow itself. With that in mind, PIV should be efficient to measure the local flow even in low SNR conditions, as it uses a cross correlation between successive frames. Nevertheless, because of the importance of Brownian motion, the cross-correlation fails at producing a peak of high SNR. Also, the geometry of the canal itself, as well as the bidirectional nature of CSF flow, makes that the use of standard large squared region-of-interest on which the cross correlation is typically calculated in PIV is not well adapted here. As particles can move in opposite directions in a 6×6 μm^2^ (32×32 pixels), the cross correlation results in several small competing peaks. Asymmetric region of interests should be more adapted to take a full advantage of the constant flow along the rostro-caudal direction, but performing a PIV with asymmetric regions is not standard and not trivial. On the other hand, PTV tracks every individual particle and should produce the trajectory of each particle versus time. One can expect that a given trajectory would then have a Brownian component, plus a directed component, each of them being separable in principle. Unfortunately here, PTV is not very robust to low SNR, and not very efficient when particle density is quite high. The most critical part is to efficiently detect the particles without error. A first solution is to have a high threshold that selects only the brightest particles, as what was done in Figure 1. Nevertheless, only a few particles could be tracked for a given dataset, and the profile could not be inferred from only a few particle positions. If the threshold is lowered, it increases the calculation time, and the probability of false detection. Since the PTV algorithm tend to minimize the distance traveled by particles, as the number of false detection increases, the false associations increases even more leading to an important underestimation of particle speed. In contrast, the automatic kymograph seemed more robust to us in this context because the flow is mostly constant in the rostro-caudal direction, which allows us to perform efficient averaging in this direction. Brownian motion is average out by averaging on 3-5 consecutive dorso-ventral positions, so that the particles produce traces long enough to be identified. The main strength of the kymograph approach is that many particles can be detected at every position, so that even if several mistakes are made, the standard error giving the error to find the correct average speed, becomes small. The longer the total acquisition time, the more particles can be averaged, and the more precise the technique is, which is not true for both PTV and PIV in which all images are independent. Nonetheless, we must admit that most of our efforts have been focused on the automatic kymograph approach, and that probably PTV and PIV algorithms may be adapted to our configuration. We could not find any available PTV and PIV algorithm that could work out directly with our data, and theories behind both techniques become more complicated in non optimal configurations. In contrast, the kymograph just requires to fit lines in an image, for which many solutions are available and just required small adjustments.

### Manual central canal diameter measurement

A Z-Stack of 30 images around the central canal of 30 hpf zebrafish embryos filled with Texas Red is acquired. A custom Matlab program is used to open, rotate, and crop the image around central canal, then to calculate the maximum intensity projection. The rotation is crucial to fit the central canal projection by a manually selected rectangle, whose width measures the canal average width. Even though the canal diameter is not constant along the rostro-caudal direction, the rectangle is placed in order to minimize the total error. The canal diameter is averaged on large rostro-caudal portions, possibly giving a fairly good estimate of its size. A random permutation of dataset order is performed to mix WT and mutants siblings, so that the manual measurement is blind from genotype and phenotype information, and to reduce potential bias. For Kurly embryos displayed in Figure 4, Texas Red was not injected, but we used the time average image of fluorescent beads filling the central canal. The results on WT siblings show remarkably similar measurements that those obtained with Texas Red, so that we decided to aggregate all these datasets together.

### Live imaging of motile cilia in central canal

30 hpf embryos from *Tg(β-actin:Arl13b-GFP; scospondin*^*icm13/+)*^ incrosses were manually dechorionated, laterally mounted in 1.5% low-melting point agarose, and paralyzed by injecting 1-2 nL of 500 µM alpha-bungarotoxin (TOCRIS) in the caudal muscles of the trunk. A spinning disk confocal microscope Leica equipped with a 40X water immersion objective (N.A. = 0.8) was used to measure cilia dynamics at a 100 frames per second for 3 seconds. Note that our estimation of high beating frequency are therefore limited to 50Hz. Because we found that the cilia average frequency in the central canal is higher than the value of 12-15 Hz previously reported^17^, we imaged cilia dynamics with 3 different microscopes (two commercial spinning disk microscopes from 3i and Leica, and one 2-photon scanning microscope using the scanning artifact, as previously described by ^7^, in order to validate our results). However, all the results displayed in this report were obtained with the same Leica commercial microscope to ensure data homogeneity. We can only postulate that the difference found in beating frequency reflects a bias for long cilia with possible slower kinematics when using differential interference contrast (DIC) to identify cilia.

### Quantification of parameters related to cilia structure and dynamics

Previously described datasets of cilia position versus time are analyzed using a custom Matlab software based on Fourier transform. A usual first step is here also to open, rotate, and crop the image around the central canal. The temporal Fourier transform of each pixel is calculated, and the position of its norm extracted and transformed into a frequency value. A 2D map of principal frequency is therefore obtained (Figure 2b2), and show distinct region of interest, possibly showing region where individual cilia can reach within one beating cycle. Within each region, the calculated main frequency can vary of a few Hz, but are grouped around a central frequency with a narrow bandwidth. From such regions, we calculated several parameters that we attribute to cilia structure and dynamics. Successive mask images of pixels with main frequency within increasing bandwidth of 5 Hz (5-10Hz, then 10 - 15 Hz, and so on until 50Hz) are created, and analyzed using Matlab image processing functions (regionprops). First, the central frequency within each region is calculated. Then, each region is associated to an ellipse, whose long axis is assumed to be cilia length. After geometrical calculation, the ellipse orientation gives the angle between the cilia and the dorso-ventral axis. The values corresponding to the negative orientations may principally originate from the few beating dorsal cilia, also oriented towards the caudal end, but from dorsal to ventral (leading to a negative angle). To account for this, we calculated the median of the absolute value of the cilia orientation in the results section. Note that it gives the same value than the median of only the positive tilted cilia (65.1° versus 65.0° for the median of absolute values). Finally, we calculate a parameter called beating height, as the cilia length times the cosine of orientation (which therefore does not depend on the sign of the orientation), which gives an estimate of which portion of central canal is occupied by beating cilia. Although some reports^43^ used the power spectral density to quantify the main beating frequency, no difference was found here between the main frequency found with the Fourier transform versus the power spectral density calculation.

### CripsR/cas9-mediated mutagenesis for *foxj1a*^*nw2*^ mutants

*foxj1a* mutants were generated upon CrispR/cas9 mediated mutagenesis as previously described^20,63^. The gRNA sequence for *Foxj1a* gene (GGACGTGTGCTGTCCTGTGC) was identified using ZiFiT Targeter website (http://zifit.partners.org/ZiFiT_Cas9) from the genomic DNA sequence obtained from Ensembl. Annealed oligos were cloned into the pDR274 plasmid (Addgene Plasmid #42250). The sgRNA was *in vitro* transcribed using MAXIscript T7 kit (Invitrogen). Cas9 mRNA was *in vitro* transcribed from pMLM3613 plasmid (Addgene Plasmid #42251) using mMessage mMachine T7 kit (Invitrogen) and polyA tailed with a polyA tailing reaction kit (Invitrogen). A mixture of 25pg of sgRNA and 600pg of Cas9 mRNA were injected in embryos at one-cell stage. CrispR/cas9-induced mutations and germ line transmission of the mutations were detected by T7EI assay and characterized by sequencing. Three separate alleles were recovered from one founder. All mutations resulted in a frame shift from amino acid 66 and an early stop codon, which was before the forkhead domain. All homozygous mutants for the allele exhibited a bent body axis. In this study, the *foxj1a*^*nw2*^ allele, which carries a 5 bp deletion, was analyzed. See more information on Supplementary Figure 3. Heterozygous adults and mutant larvae were genotyped using a KASP assay (LGC genomics) on a qPCR machine (ABI StepOne). All procedures were performed on zebrafish embryos in accordance with the European Communities Council Directive and the Norwegian Food Safety Authorities. The generation of the *Foxj1a* mutant line and fin clipping procedure on adults were approved by the Norwegian Food Safety Authorities (FOTS applications 16425 and 12395).

### Immunostaining and imaging of *foxj1a*^*nw2*^ *mutants*

For immunostaining, straight body axis and curved body axis embryos resulting from an incross of heterozygous adults were selected and processed in separate tubes. The genotype of the clutch was confirmed by KASP genotyping of siblings presenting a curved body axis. Dechorionated and euthanized embryos (collected between 26 and 30 hpf) were fixed in a solution containing 4 % paraformaldehyde solution and 1 % DMSO for at least 1 h at room temperature. Embryos were washed with 0.3 % PBSTx (3×5 min), permeabilized with acetone (100 % acetone, 10 min incubation at −20 °C), washed with 0.3 % PBSTx (3×10 min) and blocked in 0.1 % BSA/0.3 % PBSTx for 2 h. Embryos were incubated with either glutamylated tubulin (GT335, 1:400, Adipogen) or acetylated tubulin (6-11B-1, 1:1000, Sigma) [MOU1] overnight at 4 ^°^C. The next day samples were washed (0.3 % PBSTx, 3×1 h) and incubated with the secondary antibody (Alexa-labelled GAM555 plus, Thermo Scientific, 1:1,000) overnight at 4 °C. Next, the samples were incubated with 0.1 % DAPI in 0.3 % PBSTx (Life Technology, 2 h), washed (0.3 % PBSTx, 3×1 h) and transferred to a series of increasing glycerol concentrations (25 % and 50 %). Stained larvae were stored in 50 % glycerol at 4 °C and imaged using a Zeiss Examiner Z1 confocal microscope with a 20x plan NA 0.8 objective.

### Central canal photodamage and photoablation

Here, we present several experiments where we intentionally damage the central canal using different levels of light excitation. All experiments were performed after Texas Red injection (and sometimes with beads and alpha-bungarotoxin injection, but these should not play a role) by shining green (565 nm) light onto different parts of central canal. In experiments shown in Figure 4d1, a continuous medium intensity (25 mW) was applied to the entire field of view 300 μm on the left and later 300 μm on the right of the displayed field of view for 5 minutes. After a few minutes of illumination, the central canal collapses, similarly to what is found in ciliary mutants (Figure 4b2). Also, debris can be seen, as shown on the sides in Figure 4d1. Nevertheless, we never observed a breach in central canal, or fluorophores leaking out of it. A possible hypothesis is that Texas Red photobleaching due to the illumination may create damage to the central canal, or cells surrounding it, leading to an obstruction of the canal. In Figure 7 d1 and 7d2, a light pulse of 200 mW (at a wavelength of 810 nm on a two-photons microscope) was scanned 200 times on 50 pixels lined along the dorso-ventral axis, right after the end of the funnel, or at the beginning of the yolk sac extension for control siblings. This reproductively creates a hole, and hence a breach in central canal (or in the yolk sac extension tissue), around which one can observe fluorescence leaking out in the surrounding tissue.

### Bead transport analysis

To measure CSF transport, four groups of eight 30 hpf embryos (16 contracting and 16 paralyzed embryos) were injected with beads and Texas Red within a five minutes time frame. The exact injection time was saved with a resolution of one minute. Widefield fluorescence images of each embryo were acquired every 10 minutes for 60 minutes using an epifluorescence microscope (Nikon AZ100) equipped with a 2X air objective imaged the entire embryo in one image. The image creation time recorded by the computer was automatically extracted and subtracted to the injection time. The front propagation was obtained by applying a low-pass filter on all images and by manually drawing the contour from the end of the funnel until the propagation front (with a custom Matlab routine). The contour was drawn by successive lines changing direction everytime the canal changes direction. The total length was calculated as the sum of all individual length. As for other manual arbitrary measurements, a random permutation was applied to the different images to reduce bias.

### Funnel imaging and analysis

30 hpf embryos were injected with Texas Red and alpha-Bungarotoxin in diencephalic ventricle, and imaged with a confocal microscope (SP8, Leica). Because Texas Red volume is much higher in ventricles above funnel, spinning disk confocal microscope can not achieve sufficient optical sectioning to see contrast in the funnel, so that we had to use a confocal or 2 photons microscope to properly image it. For similar reason, the signal from ventricles is highly saturated to clearly visualize the funnel. A Z-stack of 50 μm around funnel position was performed and displayed as a maximal projection to visualize the entire length of the funnel and the beginning of central canal. Then, a custom Matlab routine was used to perform a manual measurement of funnel geometry. The contour of the funnel was manually drawn by drawing successive lines until the most caudal part of the image for both the dorsal and ventral walls of the funnel. We could therefore measure the local width versus the curvilinear abscissa along the line. The length of funnel was calculated between the initial point until the point of maximal diameter derivative. The width of funnel is calculated as the mean over of measured diameter on the calculated length. Note that this method is not so precise and probably overestimate the canal diameter, as proper alignment between dorsal and ventral walls is not perfect. This overestimation can be also estimated as central canal diameter is higher than when measured with previous method. Nevertheless, the ratio of about 2 between funnel and central canal diameters is supposedly and visually in the right order of magnitude.

### Full field Optical coherence tomography imaging

The full-field optical coherence tomography (FF-OCT) setup used was previously described^31,64^. The FF-OCT path is based on a Linnik interference microscope configuration illuminated by a temporally and spatially incoherent light source. A high power 660 nm LED (Thorlabs M660L3, spectral bandwidth 20 nm) provided illumination in a pseudo Köhler configuration. A 90:10 beamsplitter separates the light into sample and reference arms. Each arm contains a 40x water immersion objective (Nikon CFI APO 40x water NIR objective, 0.8 NA, 3.5 mm working distance). In the reference arm, the light is focused onto a flat silicon wafer with a reflection coefficient of about 23.5% at the interface with water. FF-OCT detects and amplifies any structure that reflects or backscatters light within the sample arm, showing a contrast based on differences in refractive index, including small extracellular vesicles with rich lipid content. The beams from sample and reference interfere only if the optical path length difference between both arms remains within the coherence length of the system, ensuring efficient optical sectioning. A 25-cm focal length achromatic doublet focuses the light to a high speed and high full well capacity CMOS camera (Adimec, custom built). The overall magnification of the FF-OCT path is 50x. The measured transverse and axial resolutions were 0.24 μm and 4 μm, respectively. Camera exposure was 9.8 ms, and images were acquired at 100 Hz. We acquired sequences of consecutive direct images and computed the standard deviation on groups of images to cancel the incoherent light that does not produce interference. Here, we also take advantage of phase fluctuations caused by the vesicle displacements occurring in the 10-100 Hz range to specifically amplify the signal while backscattered signal from the surrounding tissue is more static.

### Electron microscopy and secreted vesicles imaging

To induce expression of CD63-pHluorin, specifically labeling exosomes, embryos were injected at the 1000-cell stage. On the next day, embryos were anesthetized with tricaine and embedded in 1.5% low melting point agarose. Live imaging recordings were performed at 28°C using a Nikon TSi spinning-wide (Yokagawa CSU-W1) microscope^30^. Other embryos were used for ultrathin cryosectioning and immunogold labeling, in which case they were fixed in 2% PFA, 0.2% glutaraldehyde in 0.1M phosphate buffer pH 7.4 at 3 dpf. Zebrafish were processed for ultracryomicrotomy and immunogold labeled against GFP using PAG 10. All samples were examined with a FEI Tecnai Spirit electron microscope (FEI Company), and digital acquisitions were made with a numeric camera (Quemesa; Soft Imaging System).^30^

### Computational Fluid Dynamics (CFD) and numerical methods for solving the equations

For the numerical resolution of the system of three coupled equations detailed in the “Results” section (subsection “A bidirectional flow generated by the asymmetric distribution of beating cilia”), we used the software Matlab, from Mathworks. Our simulations were performed by the finite element method solver COMSOL, using the Creeping Flow Module. The flow simulations were carried out by solving the Stokes equation in a rectangular domain composed of 120088 domain elements and 15046 boundary elements (corresponding to 630516 degrees of freedom). The simulations were performed on a single laptop. The simulations accounting for the transport of particles in presence of a flow were performed by coupling the diffusion equation to the Stokes equation in the same rectangular domain, using the Transport of Diluted Species Module. Doubling the number of mesh elements did not affect the results obtained throughout this study, and all the simulations converged.

**The Supplementary Information includes:**

- Details on the transport model

- 4 Supplementary Figures

- 10 Supplementary Movies

**Supplementary Movie S1**. Representative example of CSF flow observed in a 30 hpf WT embryo acquired with a spinning disk microscope at 10 Hz. The video is replayed at 35 frames per second (fps). Rostral is to the left, ventral is bottom.

**Supplementary Movie S2.** Representative example of our automatic kymograph analysis of CSF flow in central canal of a 30 hpf WT embryo. For all successive dorso-ventral positions, the unprocessed kymograph (top left panel) is binarized based on intensity threshold (top right panel). Only traces with adequate length, angle, and eccentricity are kept (bottom left panel) and their orientation is measured, converted in a corresponding bead velocity, and aggregated into an histogram (bottom right panel).

**Supplementary Movie S3.** 100 Hz video of the central canal where cilia are labeled with GFP in *Tg (β-actin: Arl13b-GFP)* transgenic embryos. The video is replayed at 10 fps. Rostral is to the left, ventral is bottom.

**Supplementary Movie S4.** Representative examples of CSF flow observed in intermediate ciliary mutants (*elipsa* mutants in the two top rows, and *kurly* mutants in the bottom row), showing either purely Brownian motion, or bidirectional flow with several vortices. The raw data were acquired at 10 Hz, and the video is replayed at 20 fps. Rostral is to the left, ventral is bottom.

**Supplementary Movie S5.** 10 Hz video of 20 nm beads trapped between two ablated regions in central canal of a 30 hpf zebrafish embryo with spinning disk microscopy. The video is replayed at 35 fps. Rostral is to the left, ventral is bottom.

**Supplementary Movie S6.** One representative example of CSF flow change after contraction in the central canal, acquired at 10 Hz with spinning disk microscopy.The video is replayed at 25 fps and note that exceptionally rostral is to the right, ventral on the bottom.

**Supplementary Movie S7.** 10 Hz video of 20 nm beads in the rhombencephalic ventricle of two 30 hpf zebrafish embryos with (left panel) and without (right) heartbeat with spinning disk microscopy. The heart was stopped using butanedione. The video is replayed at 10 fps.

**Supplementary Movie S8.** 10 Hz video of 20 nm beads entering the funnel of a 30 hpf zebrafish embryo with spinning disk microscopy. The video is replayed at 35 fps.

**Supplementary Movie S9.** 10 Hz video of lateral view of a 30 hpf zebrafish embryo with full field optical coherence tomography. The video is replayed at 15 fps.

**Supplementary Movie S10.** Real-time video of central canal of a 1 dpf zebrafish embryo expressing CD63-pHluorin in the YSL.The video is replayed at 35 fps.

## Acknowledgements

We thank Prof. Brian Ciruna for the *Tg(beta-actin:Arl13b-GFP)* line, Prof. Rebecca Burdine for the *kurly* (cfap^*tm304*^ allele) mutant, Prof. Jarema Malicki for the *elipsa* mutant (*ift54/traf3ip*). We would like to acknowledge the kind and expert assistance of Aymeric Millecamps, Tudor Manoliu, and Basile Gurchenkov from the ICM.Quant imaging facility for instrument use and scientific assistance on optical imaging, as well as Sophie Nunes-Figueiredo, Bogdan Buzurin, and Monica Dicu from PHENOZFish platform for fish care. We would like to thank the Yaksi laboratory members and the fish facility support team for scientific and technical assistance. This work was funded by Human Frontier Science Program (HFSP) Research Grant (grant n° RGP063-2018), and the New York Stem Cell Foundation (NYSCF) (grant n° NYSCF-R-NI39) and an ICM postdoctoral fellowship kindly attributed to OT. The research leading to these results has also received funding from the program ‘‘Investissements d’avenir’’ ANR-10-IAIHU-06 (Big Brain Theory ICM Program), ANR-11-INBS-0011 (NeurATRIS: Translational Research Infrastructure for Biotherapies in Neurosciences), and a Helse Midt-Norge Samarbeisorganet grant to NJY and the Kavli Institute for Systems Neuroscience at NTNU.

## Author Contributions

O.T. and L.K. conceived and designed the analysis, collected and analyzed the data for the entire manuscript, with a special emphasis for biological results for O.T, and theoretical and numerical results for L.K.. Y. C.-B. helped to conceive, design analysis, and collect the data on flow experiments, cilia dynamics, and transport experiments. M.C.-T. performed the experiments on reverse transport and tail injections. N.J.-Y. generated the *foxj1a* mutant allele and characterization with immunohistochemistry. F.V. and G.V.N collected and analyzed the data concerning the fluorescence labeling of exosomes, and the electron microscopy. N.J.Y. developed and characterized the Foxj1a mutant. P.L.B. provided feedback during discussions. C.W. and F.G. conceived the project, planned experiments, modelling and analysis and provided financial support. O.T., L.K., C.W. and F.G. wrote the article with inputs from all authors.

## Declaration of Interest

The authors declare no competing interests.

## Supplementary Information Appendix

**1 FIGURE 2 - FIGURE SUPPLEMENT 1**

**FIGURE 2 - figure supplement 1.**
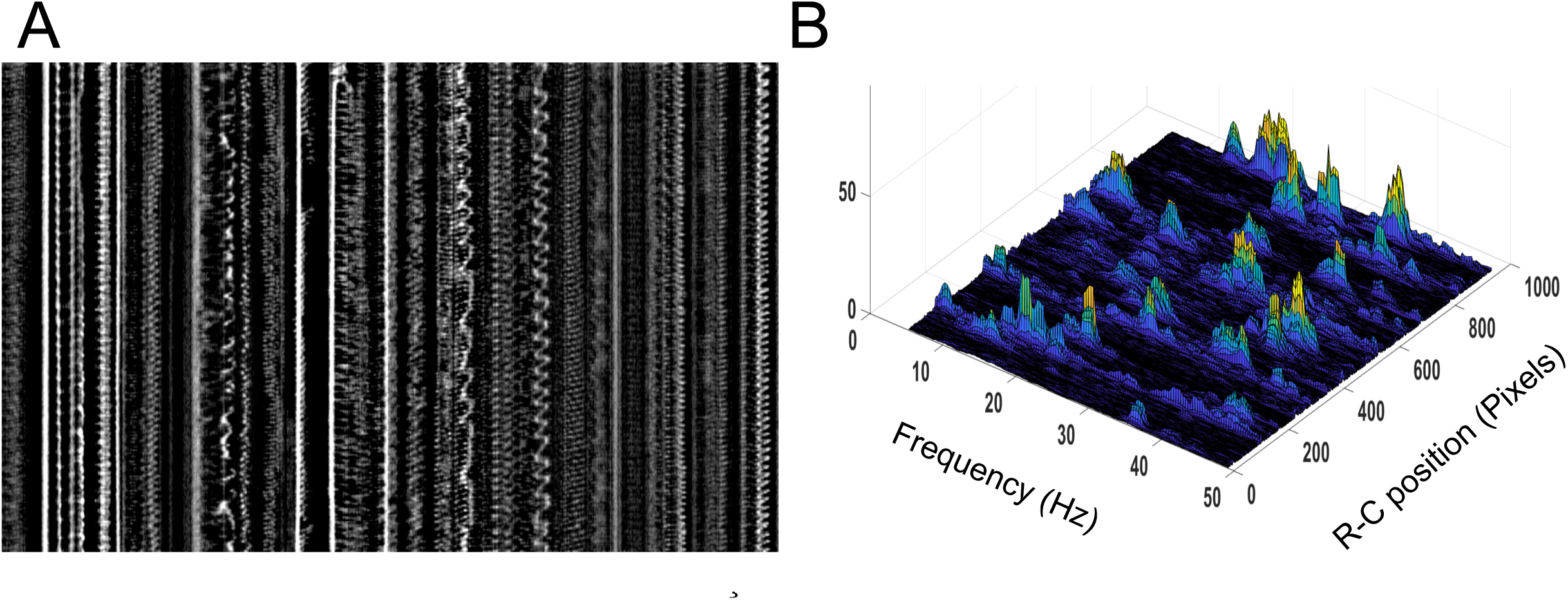
Motile cilia beating pattern in central canal. **A)** Kymograph of motile cilia dynamics in the ventral central canal associated to Figure 2B1. Note the diversity of beating patterns of neighboring cilia. Not only cilia beat at many frequencies, but some of them also show unrepeatable patterns. Horizontal scale bar is 20 *µ*m, and vertical scale bar is 0.5 second. **B)**Corresponding power spectral density map showing the position of the different maximal frequencies for each cilium. Note that some cilia show a power spectrum with several local maxima, accounting for the asynchronicity of the beating patterns observed. In order to obtain the frequency map as in Figure 2B2, we extract only the global maximum.

**II FIGURE 3 - FIGURE SUPPLEMENT 1**

**FIGURE 3 - figure supplement 1.**
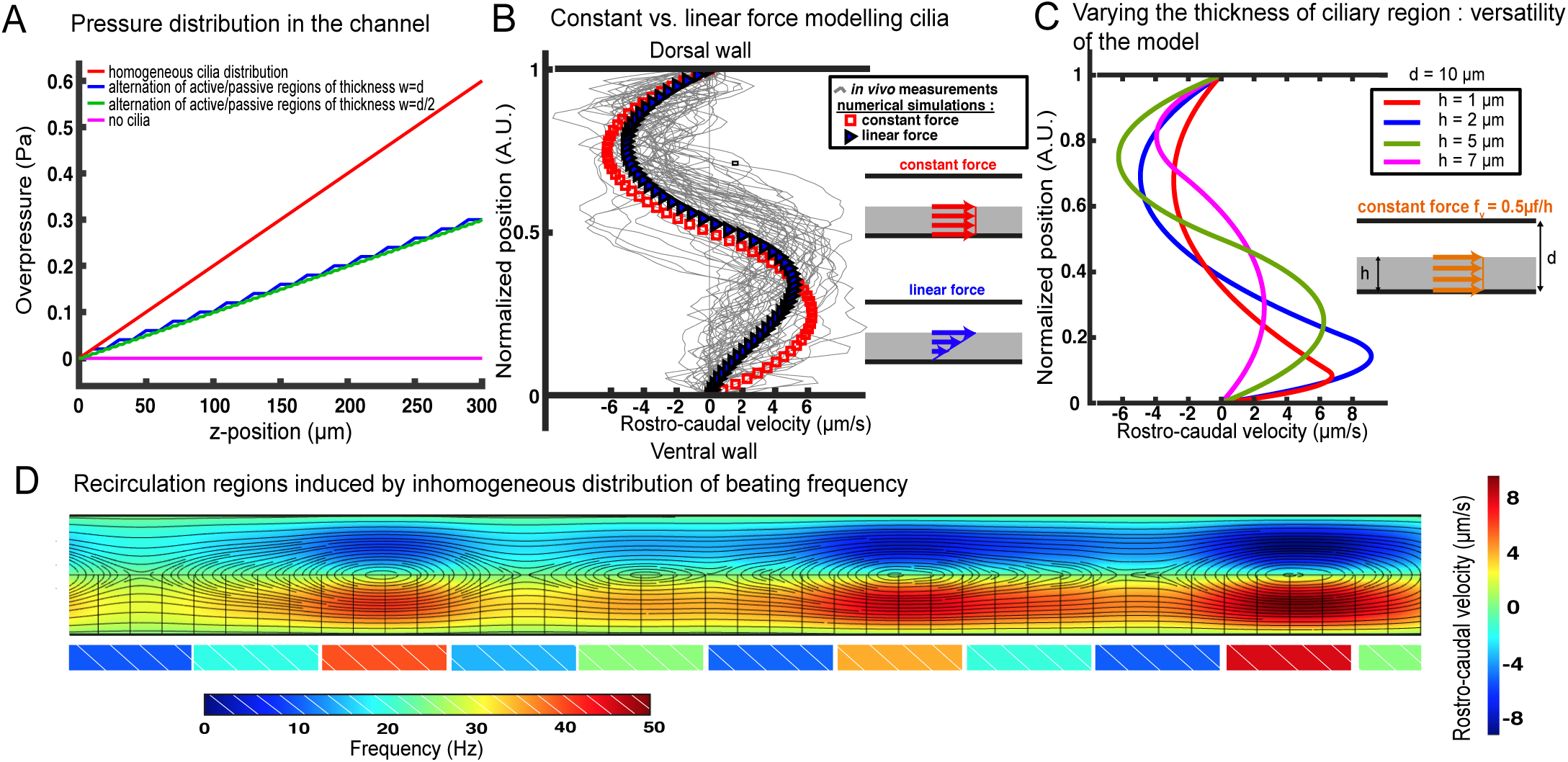
Additional information regarding our cilia-driven ow model and its consequences. **A)** Overpressure distribution in various 2-D central canals of length 300 *µ*m and diameter *d* = 10 *µ*m. The overpressure represents the excess of pressure with respect to the inlet of the CC (at the junction with the brain ventricle). In the case of a homogeneous cilia distribution (red curve), modelled as a constant force *f*_*v*_ = *µf/h*, the pressure increases linearly in the caudal direction. In the case of a loose distribution of cilia, represented by an alternation of active regions with the aforementioned force *f*_*v*_ and passive regions without any bulk force, the pressure follows a piecewise linear increase, whose frequency is controlled by the distance between active and passive regions: *w* = *d* for the blue curve and *w* = *d/*2 for the green curve. The cilia activity is thus directly correlated to the excess of pressure in the central canal, as dramatically put in evidence by the magenta flat curve in the absence of any cilia activity. This excess of pressure in the central canal induced by the cilia activity could help maintaining the lumen of the channel open, explaining why the central canal is found to be collapsed in cilia mutants (**Figure 4C**).**B)** In the main manuscript, we modelled the contribution of the cilia as a constant bulk force *f*_*v*_ = *µf/h* in the ventral region occupied by the cilia. In a prior publication, Siyahhan *et al.* (2014) modelled it as a force linearly increasing with the distance to the wall (also located in the ciliary region). We show in this figure that this choice (blue triangles) does not substantially alter the velocity profile predicted by our model (red squares), which simply turns into a 3^*rd*^ degree polynomial in the ventral half of the channel. The y-position for vanishing local velocity is slightly shifted towards the dorsal wall. **C)** Our model still holds for variations of the relative thickness of the ciliary region and the dorsal region. Following the derivation detailed in the manuscript, the velocity profiles can be obtained for arbitrary values of *h*, at fixed *t* = 10 *µ*m. The green curve corresponds to the zebrafish CC features, and the red curves corresponds to very thin cilia with respect to the diameter of the tube. This situation is representative to that encountered by Shields *et al.* (2010) with artificial magnetic cilia. They decided to model the effect of cilia as a moving wall entraining fluid. While this moving wall modelling correctly accounted for their observations in these high length ratio conditions, we point out that it would not successfully predict the flow in more common length ratios as it fully neglects the flow in the ciliary region. Conversely, our model accounts for the flow in the whole channel, which renders it very versatile to different geometries. **D)** - In **Figure 3C**, we have shown by numerical simulations that the alternation of active and passive regions can give birth to vorticity in the flow profile. Here, we show that recirculation regions can also be obtained from the succession of neighboring cilia with different beating frequencies. And the larger the difference in frequency, the larger the vortices. The black lines correspond to the trajectory of the flow, and the colormap in the channel corresponds to the rostro-caudal velocity. The colored rectangles under the ventral wall indicate the frequency for each spot in the simulation for the bulk force *f*_*v*_ = *µf/h*. This choice is consistent with the *in vivo* measurements detailed in the **Figure 2B** of the main manuscript.

**III. FIGURE 4 - FIGURE SUPPLEMENT 1**

**FIGURE 4 - figure supplement 1.**
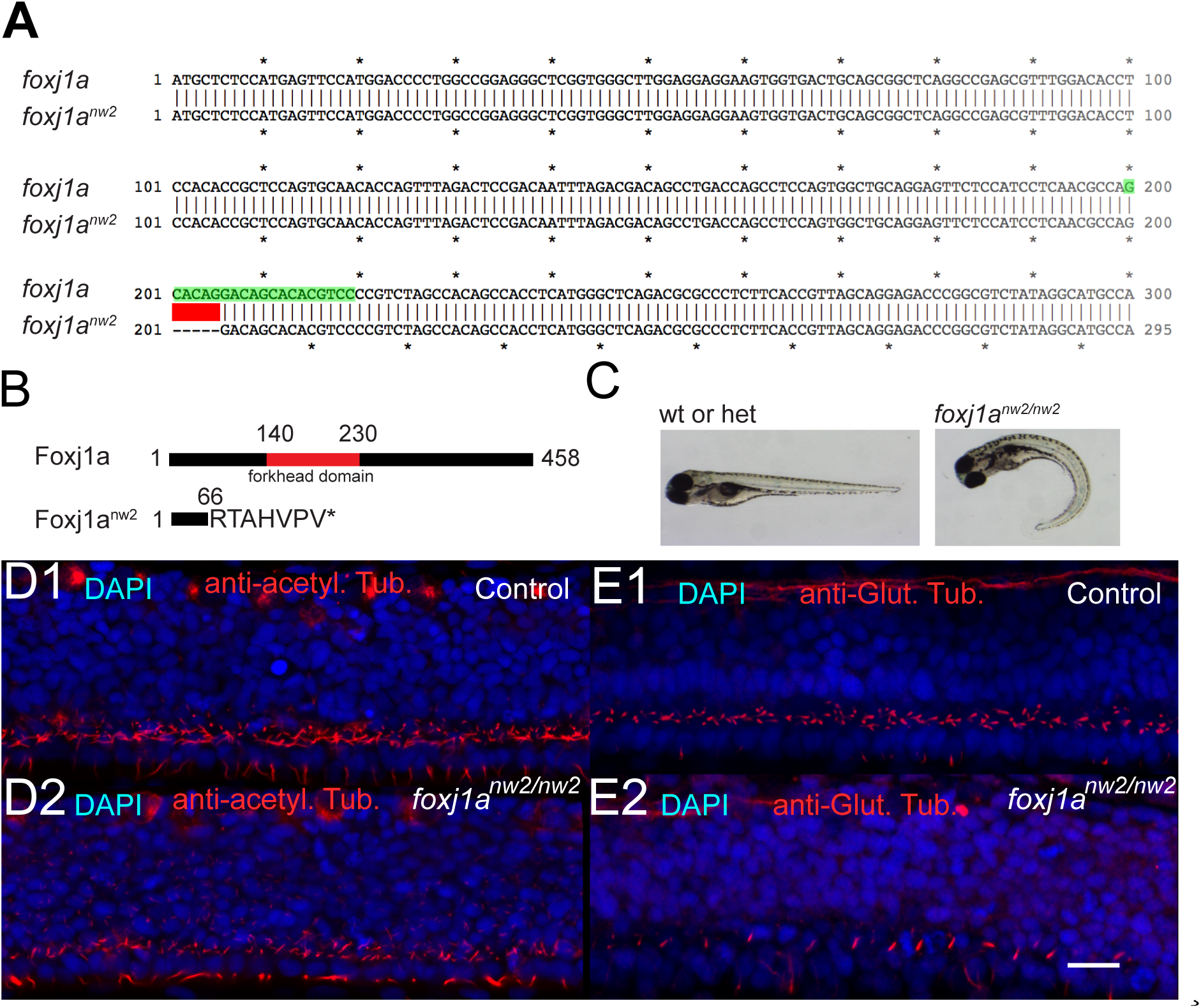
Characterization of cilia in the central canal of *foxj1a*^*nw*2^*/nw*^2^ mutant embryo. **A)** Alignment of the first 300 base pairs (bp) of the coding sequence of WT and mutant *Foxj1a*^*nw*2^ allelle reveals that the *Foxj1a*^*nw*2^ mutant allele carries a deletion of 5bp from base pair 201 to 205. The deletion is indicated in red. The gRNA sequence is highlighted in green. **B)** The Foxj1a protein consists of 458 amino acids and includes a DNA-binding forkhead domain (from amino acid 140 to 230, indicated in red). The *Foxj1a*^*nw*2^ mutant allele results in a truncated protein due to a frame shift and early stop codon and lacks the forkhead domain. **C)** - *Foxj1a*^*nw*2^*/nw*^2^ homozygous mutant larvae display a curved body axis as seen on images of a control sibling (heterozygous or WT) versus a homozygous mutant larva taken with a stereomicroscope at 4 dpf. **D)** Z-Projection stack of a lateral optical sections (Z-step = 1 *µ* over 3 sections) of the spinal cord immunostained with DAPI and against acetylated tubulin in a 30 hpf control sibling embryo (**D1**) and a foxj1a^*nw*2^*/nw*^2^ curled-down embryo (**D2**). **E)** Z-Projection stack of lateral optical sections (Z-step = 1 *µ*m over 3 sections) of the spinal cord stained with DAPI and immunostaining against glutamylated tubulin in a 30 hpf control sibling (**E1**) and a foxj1a^*nw*2^*/nw*^2^ curled-down embryo (**E2**). Scale bars are 15 *µ*m.

Both immunostaining for acetylated tubulin and glutamylated tubulin show a decrease in density, and in length of cilia in central canal in foxj1*nw*2*/nw*2 homozygous mutant embryos. Glutamylation of cilia is usually associated with active motility, which is coherent here with the position (ventral) and polarity (caudally-oriented) of glutamylated cilia in CC. The phenotype observed in the immunostaining against glutamylated tubulin usually labeling motile cilia is more drastic and suggests that most motile cilia from ependymal cells do not form in the foxj1a^*nw*2^*/nw*^2^ homogygous mutant embryos

## IV. 2D MODEL FOR THE DIFFUSIVE-CONVECTIVE TRANSPORT OF SOLUTES IN THE CENTRAL CANAL WITH A BIDIRECTIONAL FLOW-ASSOCIATED WITH FIGURE 6

We here provide further details concerning the derivation of the effective diffusivity *D*_*eff*_ of particles in the central canal in presence of a bidirectional flow.

In the CC, the motile cilia occupy the ventral half (**Fig. 2** of the main manuscript). From the flow model derived earlier (eqs. (3) and (4) of the main manuscript), one can rewrite the velocity profile changing the y-coordinate origin (*y* = 0 at the center of the channel) and using the maximal velocity *V*_*max*_ = 6 *µ*m/s:

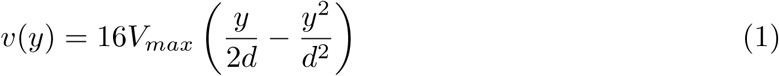

in the dorsal region (positive values of *y*), and:

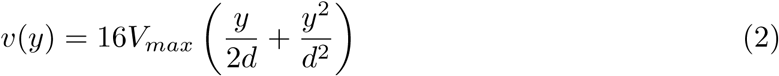

in the ventral region (negative values of *y*). The full transport equation of particles of local concentration *c* writes:

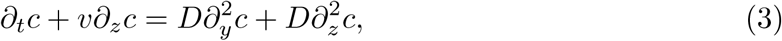

where *v*(*y*) corresponds to the two velocity profiles written previously (eqs. (1) and (2)), as the local velocity field is uniquely oriented in the rostro-caudal *z*-direction (see also **Fig. 3** of the main manuscript). D is the molecular diffusion coefficient, here taken as *D* = 10^−11^ m^2^/s, corresponding to particles of diameter 40 nm using the Stokes-Einstein law (eq. (7) in the main manuscript). Several terms of equation (3) can be neglected considering their order of magnitude. With *L* the length of the channel, and *d* its diameter, the equation (3) may be written in dimensional analysis for times of order *T* ≃ *L*^2^/*D* = *τ*_*diff*_ (characteristic spanwise diffusion time):

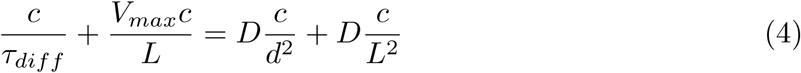

i.e.

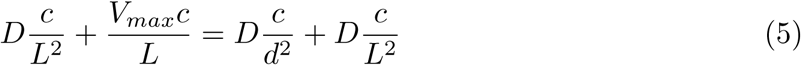

, or:

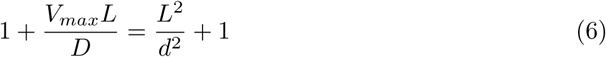

Both *V*_*max*_*L/D* and *L*^2^*/d*^2^ terms are much larger than unity, due to the high aspect ratio of the channel *L/d*. This allows to simplify equation (3) in the two dorsal (*y* > 0) and ventral (*y* < 0) regions, respectively, as follows:

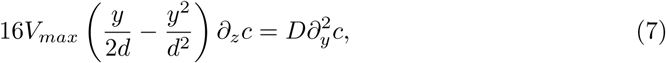

and

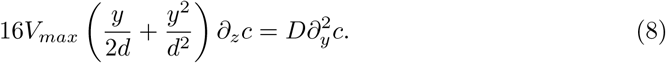

Adapting the derivation ofBruus (2008), that was done for a Poiseuille flow in a cylinder, we first assume that the axial derivative of the concentration *∂*_*z*_*c* is independent on *y*. This assumption will be verified *a posteriori* and confronted to numerical simulations. Consequently, eqs. (7) and (8) are ordinary differential equations for *c*(*y*), that can be solved. A first integration between *y* and *y* = *d/*2 for equation (7) for *y* > 0 leads to

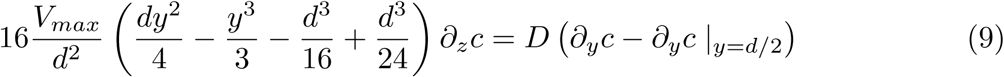

Similarly, integrating equation (8) between *y* = −*d/*2 and *y* for *y* < 0 leads to

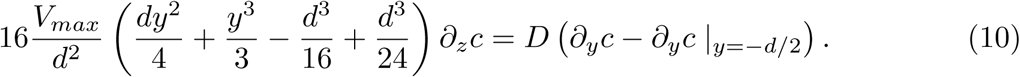

By antisymmetry, *∂*_*y*_*c* |_*y*=*d/*2_= −*∂*_*y*_*c* |_*y*=−*d/*2_. The equality of the concentration *y*-derivative in *y* = 0 then leads to *∂*_*y*_*c* |_*y*=*d/*2_= −*∂*_*y*_*c*|_*y*=−*d/*2_= 0, which simplifies equations (9) and (10). A second integration between *y* = 0 and *y* of eq. (9) for *y* > 0 leads to

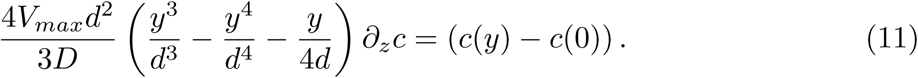

Similarly, a second integration between *y* = 0 and *y* of the equation (9) for *y* < 0 leads to

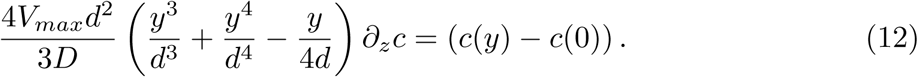

We now aim at expressing the concentration profile with respect to the average concentration 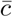 over a cross section

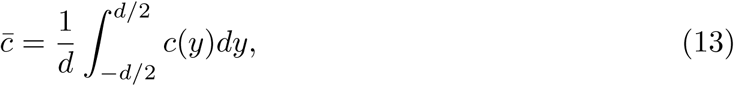

which, using eqs. (11) and (12), yields

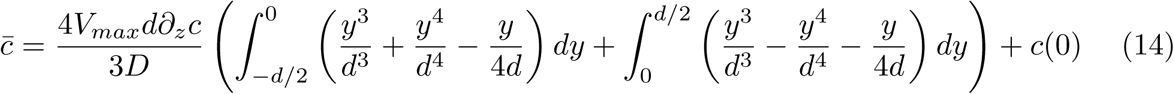

and finally

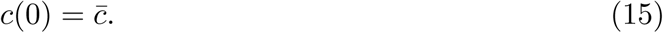

This theoretical prediction is confirmed by numerical simulations solving the complete transport equation without the assumptions made for the theoretical model (**Fig. S3A**), for different times, positions in the canal and flow velocities.

Going back to our theoretical model, one can now derive the condition for having a *y*-independent axial gradient of concentration (condition required for recovering a diffusive-like concentration profile) by differentiating the equations (11) and (12) with respect to *z*. The latter condition is consequently fulfilled only if *∂*_*z*_*c* = *∂*_*z*_*c*(0), which requires the third term to be negligible. In terms of order of magnitude, this implies that

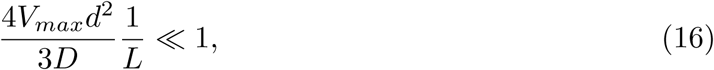

where *L* is the length of the central canal. This condition can be rewritten as a condition on the Péclet number *P é*:

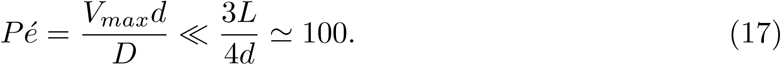

This condition on the Péclet number is always fulfilled in the CC of zebrafishes for nanobeads or lipidic vesicles (*D*∼ 10^−11^ m^2^/s), where *P é* does not exceed 10.

We now aim at deriving the effective diffusivity of particles in the CC. We calculate the average current density 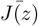 through the cross-section, using the velocity profiles (eqs. (1) and (2)) and the concentration profiles (eqs. (11) and (12)):

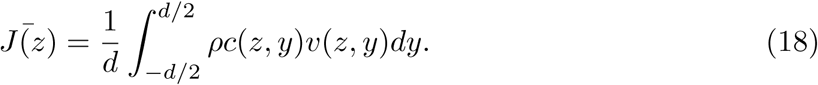

The zero average flux over the cross section 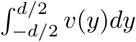 leads to the vanishing of the constant component 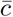 of eqs. (11) and (12). The previous expression may then be simplified:

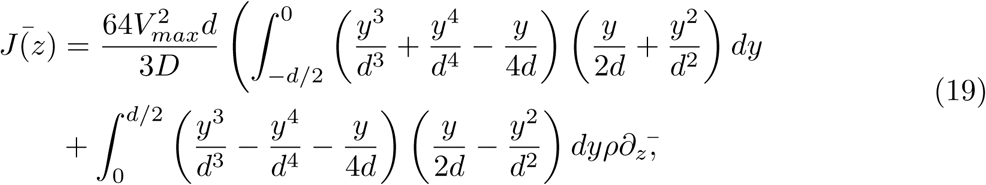

i.e.:

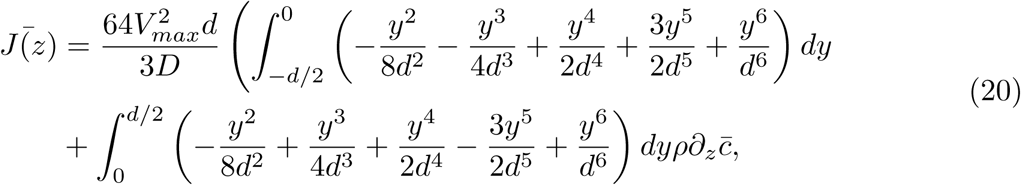

which may be approximated to

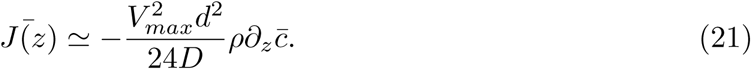

The effective diffusivity *D*_*eff*_ may then be directly from the effective Fick’s law 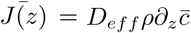:

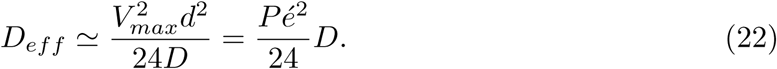

A more detailed calculation in a cylindrical channel was done by Aris (1956). It generalizes this result at intermediary Péclet numbers *P é*≃1. Adapted to our geometry and flow, it yields

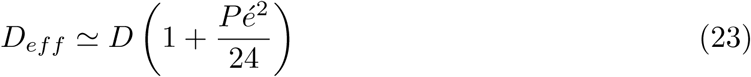

For particles of radius 20 nm (corresponding to the size of injected nanobeads), we have *D≃*5 10^−12^ m^2^/s, which leads for *V*_*max*_ = 6 *µ*m/s to an effective diffusivity *D*_*eff*_ ≃2.5*D*. Similarly, the effective diffusivity of the same particles for a faster flow of amplitude *V*_*max*_ = 20 *µ*m/s is *D*_*eff*_*≃*18*D*. These two theoretical predictions are in good agreement with the numerical simulations (**Fig. 6C** of the main manuscript).

### Relevance of the initial hypothesis

One strong assumption of the model that enabled to readily integrate eqs. (7) and (8) is that the axial derivative of the concentration *∂*_*z*_*c* is only weakly varying over a cross section, such that we consider it constant. The **Fig. S4B** plots the relative variation of the normalized concentration gradient *∂*_*z*_*c/∂*_*z*_*c*(0) as a function of *y*. It shows that this quantity exhibit minor variations of about 10% over a cross section, demonstrating the relevance of the simplifying assumption.

**V. FIGURE 6 - FIGURE SUPPLEMENT 1**

**FIGURE 6 - figure supplement 1.**
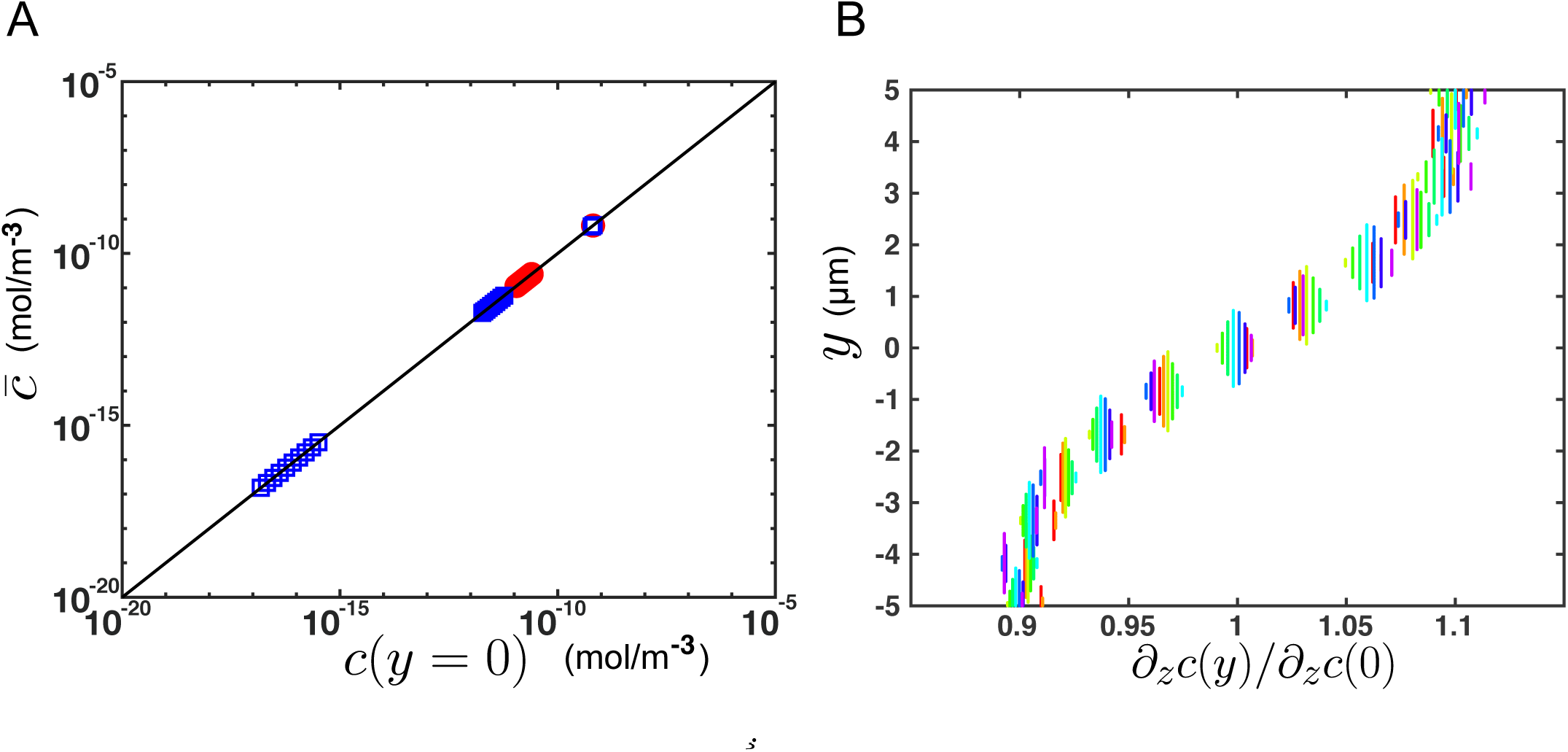
Additional results from numerical simulations supporting the theoretical model for transport. **A)** Average concentration over a cross section versus concentration at the center of the CC (*y* = 0), for different flow velocities (*V*_*max*_ = 6 or 20 *µ*m/s), at different times and for different *z*-position in the channel, obtained from numerical simulations (FEM, COMSOL) on a 2-D central canal. The black line corresponds to 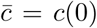 and confirms the predictions of the theoretical model. **B)** - Relative variation of the axial gradient of concentration *∂*_*z*_*c* over a cross section (*y*-position), at various *z*-positions (corresponding to different colors), from numerical simulations. *∂*_*z*_*c* varies of about 10% around its mean value over a cross section, which justifies the assumptions done in the model (from eqs. (7) and (8) to eqs. (9) and (10)) to neglect its variations in comparison with the strong variations of the other integrand *y*^3^*/d*^3^ + *y*^4^*/d*^4^ − *y/*4*d*.

